# Profiling Mouse Brown and White Adipocytes to Identify Metabolically Relevant Small ORFs and Functional Microproteins

**DOI:** 10.1101/2022.03.12.484025

**Authors:** Thomas F. Martinez, Sally Lyons-Abbott, Angie L. Bookout, Cynthia Donaldson, Joan M. Vaughan, Calvin Lau, Ariel Abramov, Arian F. Baquero, Karalee Baquero, Dave Friedrich, Justin Huard, Ray Davis, Bong Kim, Ty Koch, Aaron J. Mercer, Ayesha Misquith, Sara A. Murray, Sakara Perry, Lindsay K. Pino, Christina Sanford, Alex Simon, Yu Zhang, Garrett Zipp, Maxim N. Shokhirev, Andrew J. Whittle, Brian C. Searle, Michael J. MacCoss, Alan Saghatelian, Christopher A. Barnes

## Abstract

The absence of thousands of recently annotated small open reading frame (smORF)-encoded peptides and small proteins (microproteins) from databases has precluded their analysis in metabolism and metabolic disease. Given the outsized importance of small proteins and peptides such as insulin, leptin, amylin, glucagon, and glucagon-like peptide-1 (GLP-1) in metabolism, microproteins are a potentially rich source of uncharacterized metabolic regulators. Here, we annotate smORFs in primary differentiated brown, white, and beige mouse adipose cells. Ribosome profiling (Ribo-Seq) detected a total of 3,877 unannotated smORFs. Analysis of RNA-Seq datasets revealed diet-regulated smORF expression in adipose tissues, and validated the adipose translation of the feeding-neuron marker gene Gm8773. Gm8773 encodes the mouse homolog of FAM237B, a neurosecretory protein that stimulates food intake and promotes weight gain in chickens. Testing of recombinant mFAM237B produced similar orexigenic activity in mice further supporting a role for FAM237B as a metabolic regulator and potentially part of the brain-adipose axis. Furthermore, we demonstrated that data independent acquisition mass spectrometry (DIA-MS) proteomics can provide a sensitive, flexible, and quantitative platform for identifying microproteins by mass spectrometry. Using this system led to the detection of 58 microproteins from cell culture and an additional 33 from mouse plasma. The proteomics data established the anti-inflammatory microprotein AW112010 as a circulating factor, and found that plasma levels of a microprotein translated from a FRS2 uORF is elevated in older obese mice. Together, the data highlight the value of this database in examining understudied smORFs and microproteins in metabolic research and identifying additional regulators of metabolism.

## INTRODUCTION

Recent advances in ribosome profiling (Ribo-Seq) and proteogenomics have identified thousands of unannotated peptides and small proteins (microproteins or MPs, less than 150 amino acids) encoded from small open reading frames (smORFs) in mammalian genomes (Chen et al., 2020; Ingolia et al., 2011; Martinez et al., 2020; Slavoff et al., 2013). Several of these newly annotated smORFs have been characterized with biologically significant activities ranging from cell biology to physiology. For instance, there are microproteins involved in cell stress and survival (Chen et al., 2020; Chu et al., 2019) and several microproteins that have roles in muscle development and function (Anderson et al., 2015; Bi et al., 2017; Nelson et al., 2016; Zhang et al., 2017). The application of CRISPR screening with single cell transcriptomics has enabled unbiased genetic screens to identify novel functional smORFs (Chen et al., 2020). Together, these studies reveal a large group of unexplored mammalian genes with functions in biology and even pathology. Indeed, gene therapy with the DWORF microprotein mitigates cardiomyopathy in mice (Makarewich et al., 2018).

With a current growing population of over 600 million obese adults, the health impacts of obesity are numerous and still unfolding (The GBD 2015 Obesity Collaborators, 2017). The underlying causes of obesity are multifactorial with individual genetics, nutritional sources, and physical activity all playing roles. Where fat is deposited and processed organism-wide, along with the genomic and molecular underpinnings of the differing fat depots and their distributions and functions, are still being studied. A survey of biology reveals that peptides and small proteins have an outsized role in regulating metabolism, suggesting that microproteins might be an unusually rich source of metabolic regulators. In support of this hypothesis, several microproteins have recently been discovered with roles in cellular metabolism (fatty acid oxidation, respiration), such as the 56-amino acid microprotein from LINC00116 mRNA that is now called mitoregulin (Chugunova et al., 2019; Friesen et al., 2020; Lin et al., 2019; Makarewich et al., 2018; Stein et al., 2018). Additionally, a microprotein derived from mitochondrial RNA called MOTS-c is a circulating factor that reduces obesity and ameliorates insulin resistance (Lee et al., 2015). smORF discovery is still early and additional profiling experiments in new cells and tissues continue to identify novel potentially microprotein-coding genes (Chen et al., 2020; Martinez et al., 2020; Prensner et al., 2021). Therefore, we reasoned that a search for additional metabolically important smORFs should focus on the unexplored adipose tissue smORFome.

Adipose tissue is critical in health and disease, and functions as a physiological energy depot as well as an endocrine tissue that is able to secrete peptide and protein factors that regulate feeding, energy balance, and thermogenesis to varying degrees depending on the species (Kusminski et al., 2016; Reilly and Saltiel, 2017; Smith and Kahn, 2016). We examined primary mouse differentiated brown, subcutaneous white, and beige (differentiated from subcutaneous white) adipocytes with Ribo-Seq to generate a database of 3,877 unannotated smORFs in these cells. Here, we underscore the utility of this database as a resource for the metabolism community by demonstrating how this data can be used to identify potentially translated smORFs with Ribo-Seq and their associated microproteins with mass spectrometry (MS). These smORF-encoded microproteins could be important regulators of metabolism, as we show a circulating microprotein with an obesity- and age-related level increase as well as the validation of the translation and functional characterization of a mouse microprotein from the predicted gene Gm8773 mRNA, that controls feeding when administered centrally to diet-induced obese mice.

## RESULTS

### Ribo-Seq identification of microprotein-coding smORFs in primary mouse brown, white, and beige fat cells

Adipose tissue is a complex mixture of cell types that includes adipocytes, fibroblasts, vascular cells, and immune cells (Vijay et al., 2020). Using primary mouse adipocytes focused the discovery efforts on adipocyte-specific microprotein-coding smORFs without contamination from other cell types. Previous studies in human cell lines demonstrated that while some smORFs are translated in multiple cell lines, many translated smORFs are cell line specific (Martinez et al., 2020). Therefore, comprehensive profiling of adipose-derived smORFs requires annotation of different adipose cell types. The primary types of adipose tissue are thermogenic brown adipose tissue (BAT) and energy storage-focused white adipose tissue (WAT) with WAT being subdivided into subcutaneous and visceral depots. Additionally, the subcutaneous WAT (scWAT) depot has thermogenic potential in the “beige” phenotype, so we sought to annotate translated smORFs in primary cells derived from BAT and from scWAT differentiated into both white and beige phenotypes separately (Cannon and Nedergaard, 2004; Rosen and Spiegelman, 2014).

Primary brown adipocytes and subcutaneous white adipocytes were prepared from freshly dissected BAT and scWAT, respectively, isolated from 7 week old female C57Bl6/J mice and differentiated using established protocols (Cannon and Nedergaard, 2001; Hausman et al., 2008) (Fig. 1A). Additionally, white adipocytes were also grown under beiging conditions (Roberts et al., 2014). Promoting brown and beige adipocyte biogenesis is a potential therapeutic approach for increasing energy expenditure to reduce obesity (Kajimura, 2015), which was the rationale for also analyzing beige subcutaneous adipocytes.

**Fig. 1.**
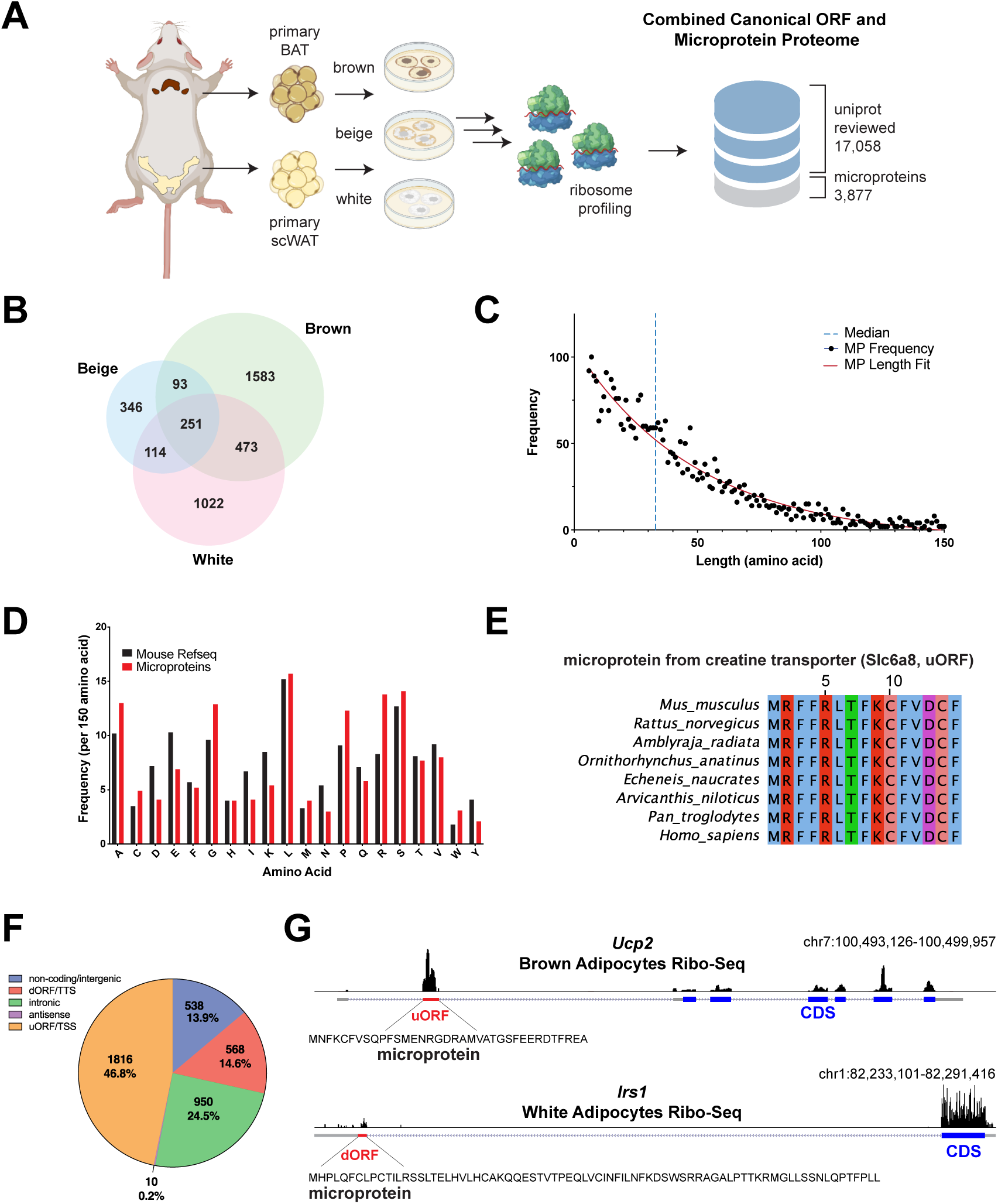
Ribosome profiling to define primary differentiated white, brown, and beige mouse adipocyte microproteins. (A) Primary subcutaneous white, beige (from subcutaneous WAT), and brown adipocytes were generated from freshly isolated subcutaneous white adipose tissue (WAT) and brown adipose tissue (BAT). Primary differentiated adipocytes were then analyzed by Ribosome Profiling (Ribo-Seq), leading to the identification of thousands of microprotein-encoding small open reading frames (smORFs) not found in curated databases (UniProt, Refseq, Ensembl). (B) A Venn Diagram of the identified smORF distributions between primary white, beige, and brown adipocytes. (C) The length distribution of the microproteins encoded by smORFs results in a median length of 33-amino acids (blue dotted line). (D) Comparison of the amino acid composition of microproteins (red bars) to mouse RefSeq proteins (black bars). (E) Homer analysis of the microprotein-encoding smORF positions was used to estimate numbers of smORFs in non-coding/intergenic regions, those downstream of a CDS (downstream ORF (dORF)/translational termination site (TTS)), intronic regions, antisense RNAs, and upstream of a CDS (upstream ORF (uORF)/translational start site (TSS)). (F) Primary differentiated brown adipocyte Ribo-Seq data shows ribosome occupancy on a uORF of the *Ucp2* gene, and subcutaneous white adipocyte Ribo-Seq data detected a dORF in what was considered the 3’-UTR of *Irs1*. (G) Some microproteins such as the 14-amino acid microprotein encoded from the creatine transporter Slc6a8 uORF are evolutionarily conserved.

We used a combination of Ribo-Seq and RNA-Seq to comprehensively annotate protein-coding smORFs in these cell lines (Martinez et al., 2020). Ribo-Seq is a powerful tool for the unbiased identification of translated open reading frames (ORFs) that relies on footprinting of ribosomes and sequencing of ribosome protected RNA fragments (RPFs) to determine the transcriptome-wide location of ribosomes (Ingolia et al., 2009, 2011). After sequencing, RPFs are mapped onto a *de novo* assembled transcriptome where RNA-Seq defined transcripts from the same cell type are used to create a database of all possible ORFs by in silico three-frame translation. Each possible ORF, including smORFs, is then scored for likelihood of translation using the RibORF computational pipeline (Ji, 2018; Ji et al., 2015) which defines protein-coding ORFs undergoing translation. The final list of unannotated predicted protein-coding smORFs is assembled by removing ORFs that are annotated in the SwissProt database as reviewed protein-coding ORFs in UniProt as well as smORFs that contain regions of overlap with annotated ORFs in the UCSC mouse gene database, where the overlap creates difficulty in accurately assessing translation of the smORF versus the canonical ORF using Ribo-Seq. We previously showed that Ribo-Seq identification of smORFs is inherently noisy, most likely because smORF RNAs are on average at lower concentrations than those of annotated ORFs (Martinez et al., 2020). To improve confidence in smORFs called translated, we collect multiple replicates for each sample (Table S1).

Ribo-Seq analysis for all three cell types identified a total of 3,877 previously unannotated smORFs (Fig. 1A and Table S1). Of these, 251 smORFs were called translated in all three cell types, while 931 were identified in at least two cell types (Fig. 1B). These results are consistent with number of predicted protein-coding smORFs identified previously in a variety of human cell lines (Chen et al., 2020; Martinez et al., 2020). The length distribution of mouse adipose microproteins also follows a similar curve as human microproteins and the 33 amino acid median length is close to the 32 amino acid median identified for novel human microproteins (Martinez et al., 2020) (Fig. 1C). Next, an analysis of the amino acid frequencies in predicted mouse adipose microproteins showed higher usages of alanine, glycine, proline, arginine, and tryptophan as well as lower levels of aspartate, glutamate, asparagine, glutamine, and tyrosine, which is identical to what we had observed in human microproteins (Martinez et al., 2020) (Fig. 1D). We also analyzed the microproteins for evidence of transmembrane domains using the TMHMM Server (Sonnhammer et al., 1998) and signal peptides using the SignalP and Phobius analysis tools (Almagro Armenteros et al., 2019; Käll et al., 2004), and found that only 2.8% contain at least one predicted transmembrane helix and 5.5% contain a predicted signal peptide secretion tag, similar percentages to what is observed with human microproteins (Table S2).

In addition, 204 predicted protein-coding smORFs have a positive average PhyloCSF score, suggesting higher likelihood of being a conserved coding region (Table S1). When employing tBLASTn (Gertz et al., 2006), we also observed 241 smORFs with high amino acid sequence similarity to ORFs found on human transcripts, which indicates possible conservation between mouse and human. For example, the microprotein encoded by a smORF in the 5’-UTR of the creatinine transporter (Slc6a8) is identical in several species (Fig. 1E). Microprotein sequence conservation is often observed in functionally characterized microproteins, and thus suggests that this set of microproteins are the most likely to produce functional peptides or small proteins. However, there are examples of functional microproteins that are not well conserved, such as the mouse AW112010 microprotein which is involved in mouse mucosal immunity but is not conserved in humans (Jackson et al., 2018). In aggregate, the data indicate smORFs are potentially translated to a similar extent across different mammalian species, and share many of the same global properties. Furthermore, as in humans, many of the microproteins identified in mouse adipocytes are likely to be functional based on their conservation (Chen et al., 2020; Martinez et al., 2020).

### smORFs are frequently located on genes important in lipid metabolism

The majority of predicted protein-coding smORFs in mouse adipocytes overlap with or are within close proximity to annotated RefSeq transcripts. Representing the largest category, 1,816 smORFs (∼50%) are found upstream of annotated coding sequences within or proximal to 5’-untranslated regions (Fig. 1F). The existence of upstream ORFs (“uORFs”) has been known for decades (Andreadis et al., 1982; Rose and Botstein, 1983) and experiments on the yeast GCN4 (Miller and Hinnebusch, 1990) and mammalian ATF4 (Lee et al., 2009) have demonstrated a role for these ORFs in the cis regulation of downstream translation (Kozak, 2002). The advent of genomics technologies including Ribo-Seq demonstrates that uORFs are prevalent (Calvo et al., 2009), with Ribo-Seq confirming their potential translation (Chen et al., 2020; Ingolia et al., 2009, 2011; Martinez et al., 2020).

There are reports of uORFs mediating the translation of metabolically important genes. To determine whether translated uORFs identified in our data could potentially regulate metabolic pathways, we employed gene ontology (GO) analysis on the annotated genes containing uORFs (Ashburner et al., 2000). The top 10 redundancy-filtered pathways regulated by genes containing uORFs included lipid metabolic processes, positive regulation of catabolic processes, mitochondrion organization, and glycoprotein metabolic process (Fig. S1 and Table S3). Of note, our Ribo-Seq data detects uORFs on genes known to be involved in metabolism, including *Insr, Igf1, Cd36, Adra1a, Adrb1, Ppara, Plin1, Irs2, Ucp1*, and *Ucp2* (Fig. 1F). The biological significance of most of these uORFs is unknown, but the *Cd36* uORFs were previously characterized and are necessary for glucose regulation of *Cd36* translation (Griffin et al., 2001). The translation of each of these smORFs and others in the data, might regulate the translation of these canonical ORF genes critical to metabolism. It is also possible that microproteins encoded on uORFs could have functions independent of translational regulation of the canonical coding ORF (Chen et al., 2020). For instance, a uORF on *Pten* encodes a 31-amino acid microprotein that regulates lactate metabolism in cells in a mouse glioblastoma model (Huang et al., 2021) and a recent report shows a uORF-encoded peptide that acts as an autoinhibitor of PKC signaling (Jayaram et al., 2021). Taken together, it is evident that understanding of the functional significance of uORFs in biology is still unfolding. The potential for each uORF to either be involved in translational control of the downstream ORF via ribosome stalling or be translated into a functional stable microprotein with functions on its own, or even both, is unclear. Evolutionarily, it has been shown that mutations to the stop codon of uORFs are under a stronger negative selective pressure than missense mutations in the main ORF corroborating their importance in biology irrespective of the mechanism (Lee et al., 2021).

Beyond uORFs, another 568 smORFs (∼15%) are located downstream of annotated coding sequences within or proximal to 3’-untranslated regions (Fig. 1E). In contrast with uORFs, these downstream open readings frames (dORFs) have been shown in some instances to enhance translation of the gene’s main coding ORF (Wu et al., 2020). For instance, we find a prominent dORF on the *Irs1* mRNA that suggests the potential for translation level regulation of the IRS1 protein (Fig. 1G). An additional 950 smORFs (∼25%) overlap annotated intronic regions (Fig. 1F). Finally, 538 smORFs (∼10%) are located on either annotated non-coding RNAs or within intergenic regions, both of which represent transcripts that lack any annotated coding regions (Fig. 1F). Moreover, intergenic smORFs are encoded on transcripts that are entirely unannotated in RefSeq. In total, we observed that protein-coding smORFs in mice are found throughout the transcriptome in regions previously considered untranslated, which is consistent with previously published Ribo-Seq based smORF discovery in other cell and tissue types.

### Data-independent acquisition mass spectrometry (DIA-MS) allows for proteome-wide quantification of microproteins with single-peptide resolution

Only a fraction of smORFs produce functional peptides and microproteins. MS-based proteomics complements Ribo-Seq by validating translation and demonstrating that the translated microprotein is stable and long lived enough for detection. Nevertheless, detecting microproteins by MS-based proteomics is challenging because most peptides and microproteins are at low concentrations while also generating few detectable peptides. Consequently, almost all microproteins are identified by detecting a single peptide. Common MS methods such as data-dependent acquisition (DDA) stochastically sample the proteome, leading to individual peptides being analytically missed across biological replicates (Gillet et al.; Ludwig et al., 2018; Navarro et al., 2016; Pino et al., 2020; Searle et al., 2018; Ting et al., 2017), creating a problem for quantifying microproteins that liberate few, if any, peptides (Fernandez-Costa et al., 2020; Ma et al., 2016). To circumvent these issues and improve our chances for quantifying microproteins, we chose a DIA-MS method that favors the reliable quantification of individual peptides, while also being fast enough for proteome-wide quantitation (Fig. 2) (Searle et al., 2018). This approach uses a pool of sub-aliquots from all sample groups (depicted as blue/red) to generate a sample-specific “chromatogram library”. Here the pool is analyzed using both column fractionation followed by DDA and gas-phase fractionated (GPF) DIA. Peptides are identified from the DDA injections, and the DIA-specific peak shape, retention time, and fragmentation patterns for those peptides are assigned from the GPF-DIA injections. These peptide parameters are compiled in the chromatogram library and used to accurately detect and quantify peptides in individual samples.

**Fig. 2.**
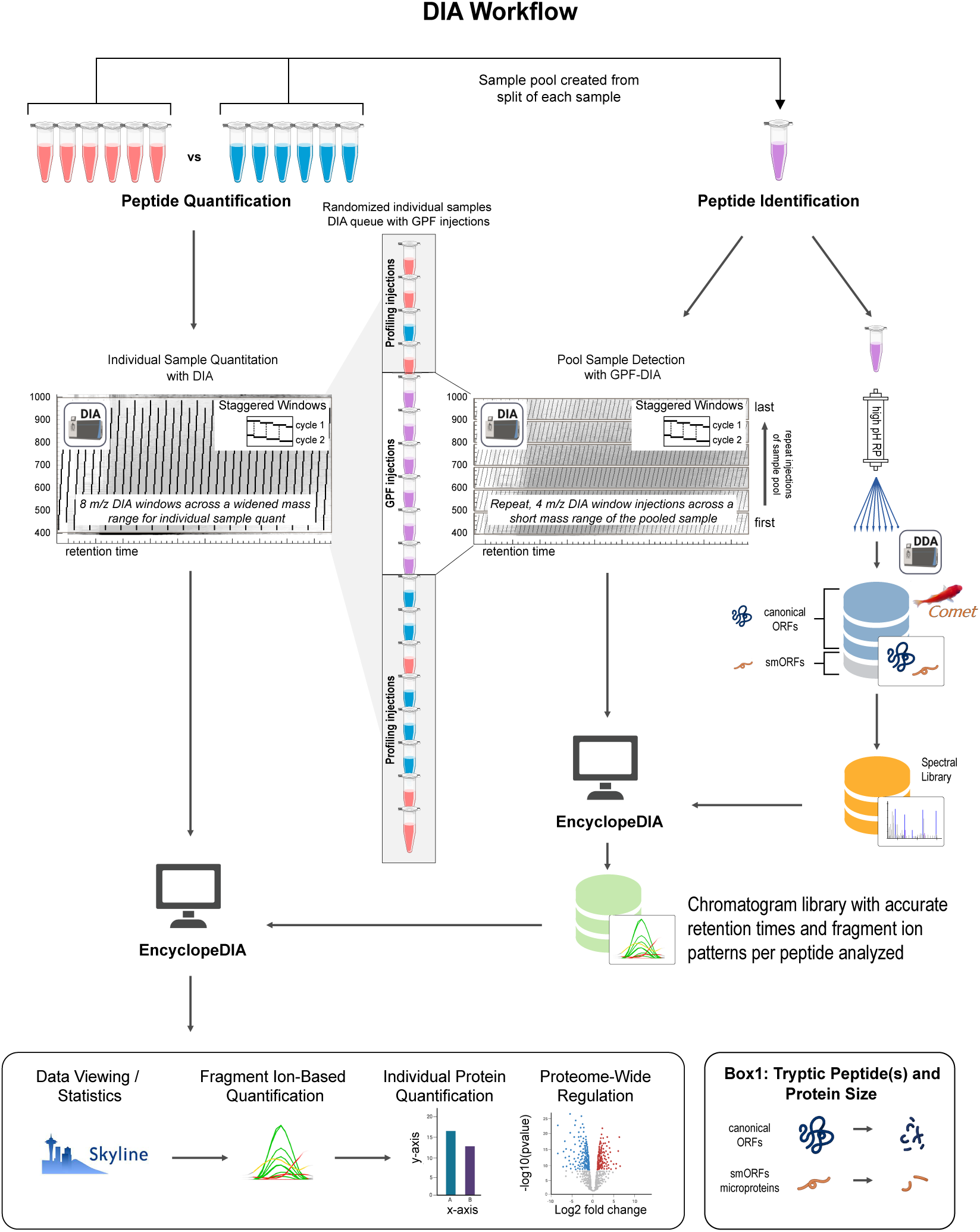
Schematic of Data-independent Acquisition Mass Spectrometry (DIA-MS) method that employs the chromatogram library approach. An idealized experiment with two groups of six (red and blue) is depicted in the upper left. A small volume of each sample is split and pooled for peptide identification analysis (purple). First, the pool is subjected to high pH reversed-phase (RP) fractionation where peptide identifications are generated from DDA injections of each RP fraction. The RiboSeq generated smORF-encoded microprotein sequences are amended to the canonical reviewed UniProt proteome database (blue/grey library icon) where each RP fraction DDA injection is searched (Comet + PeptideProphet) with the results used to make a spectral library of peptide identifications (orange library icon). The pool is also subjected to a small (4 m/z) DIA gas-phase fractionation (GPF) window. The GPF chromatogram library method utilizes 6 replicate injections of the sample pool with small, staggered windows (4 m/z) across a short mass range (100 m/z). Each short mass range injection covers one sixth of the total mass range of the profiling method used on each sample. The profiling method for quantifying each sample uses larger staggered windows (8 m/z) from a mass range of 400-1000 that covers the entire mass range of the GPF injections. The GPF injections are inserted into the randomized queue of individual samples to be quantified, allowing very accurate retention time realignment and improved accuracy in the peptide extractions from the DIA-MS profiling data. EncyclopeDIA is first used to create the chromatogram library (green library icon) that contains all of the DIA-based peptide identifications and accurate retention times. EncyclopeDIA is then used again to extract fragment ion-based peptide information from the profiling injections using the chromatogram library generated from the GPF injections. Extracted ion chromatograms for each peptide are then viewable in Skyline (MacLean et al., 2010), allowing for viewing the data, proteome regulation analysis (generation of protein or peptide volcano plots), and exporting the summed peptide fragment ion intensities for each peptide. Multiple peptides per protein can be summed to quantify individual proteins, but the method is also highly accurate on a single peptide level. Box 1 depicts how this methodology is important for smORF-encoded microprotein discovery. Each smORF sequence may only generate a small number of analytical peptides in a tryptic digest (or any given sample prep methodology). In contrast, a canonical ORF may generate many analytical peptides per protein in a tryptic digest.

### Microprotein proteomics with DIA-MS in primary differentiated brown and subcutaneous white adipocytes confirms novel translation and imparts evidence for secretion

We used DIA-MS to validate the translation of smORFs and to quantify differences in microprotein levels between whole cell lysates and the secreted protein space (“secretomes”) of primary differentiated BAT- and scWAT-derived cells (Fig. 3A). BAT-derived, differentiated “brown” cells have distinctly smaller lipid droplets compared to the differentiated scWAT-derived “white” cells (Fig. 3A). We validated this technique by examining known protein changes between brown and white cells. Mitochondrial brown fat uncoupling protein 1 (UCP1) was characteristically elevated in the brown cultures (Fig. 3B). Both types of adipocytes produced perilipin 1 (PLIN1), which is the major packaging protein for lipid droplets, and, as expected, we observed higher levels in BAT-derived brown cultures with their numerous smaller lipid droplets. Additionally, UCP1 (Cannon and Nedergaard, 2004) and glucose transporter type 4 (GLUT4) (Orava et al., 2011; Shimizu et al., 1993) protein levels were higher in brown adipocytes, consistent with the literature (Fig. 3B). Analysis of the secretome was also consistent with expectations as we observed no difference in adiponectin (ADIPOQ) levels between cell types but higher apolipoprotein E (APOE) amounts in the brown adipocytes culture media (Bartelt et al., 2017). Taken together, these results from the comparison of primary differentiated brown and subcutaneous white adipocytes (BAT-and scWAT-derived, respectively) demonstrate that our cultures are generating functional surrogates of their respective tissue depots, and that our quantitative proteomics methods are accurately detecting significant biological differences.

**Fig. 3.**
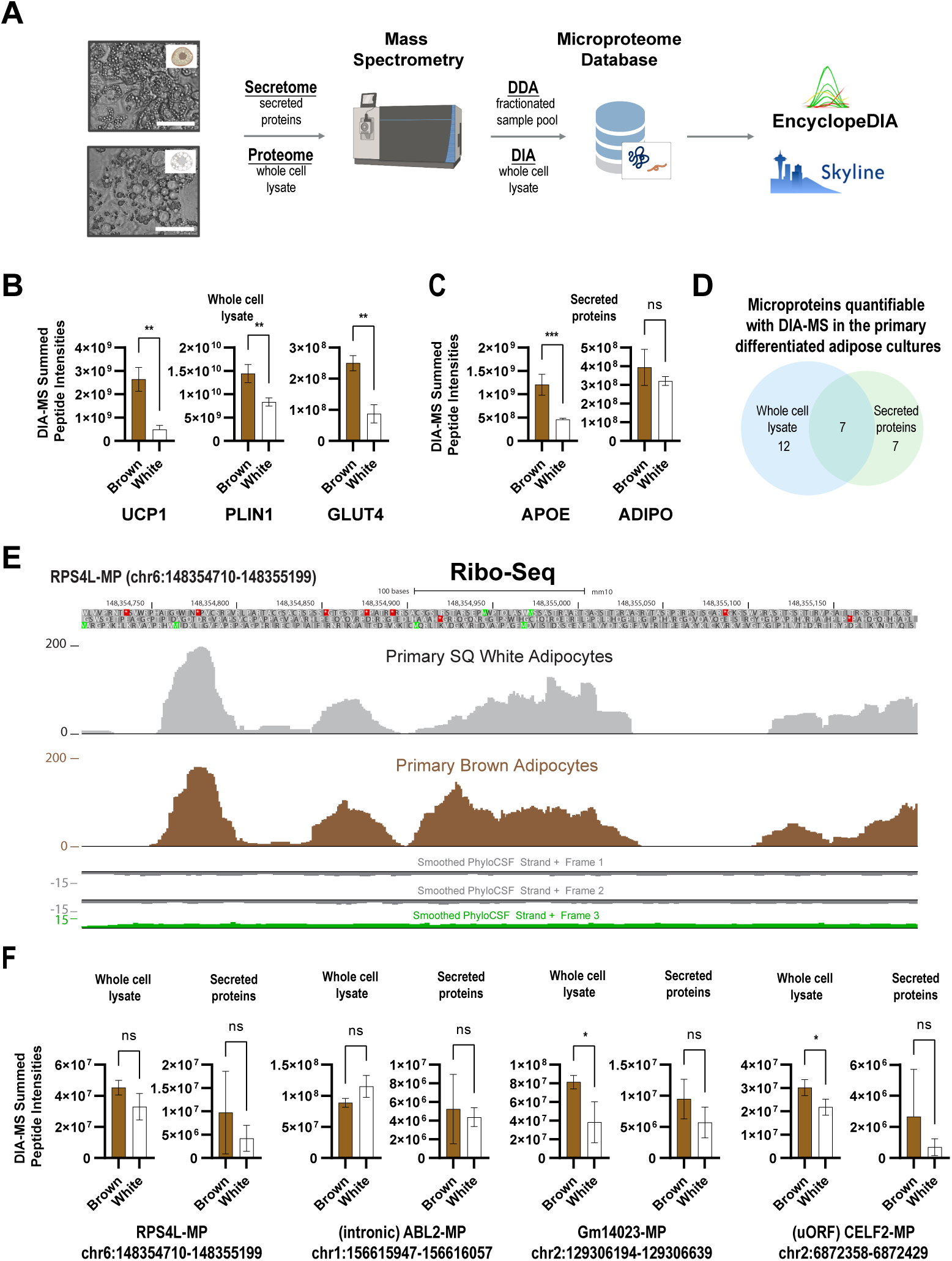
DIA-MS quantitation of canonical ORF proteins and small ORF microproteins in primary differentiated brown and subcutaneous white adipocytes. (A) A schematic of the experimental design quantifies the proteomes of primary differentiated brown and subcutaneous white adipocytes, allowing for identification with DDA and quantification with DIA-MS of canonical ORF proteins smORF-encoded microproteins in whole-cell lysates and conditioned media (secretomes). (B) DIA-MS quantification of canonical ORF proteins UCP1, PLIN1, and PCK1 from the whole-cell lysates of the same primary differentiated brown and subcutaneous white adipocytes analyzed with RiboSeq in Figure 1. (C) DIA-MS quantification of known secreted proteins APOE and ADIPO (Adiponectin). (D) Venn diagram of the total number of microproteins quantified with DIA-MS in the whole-cell lysates and conditioned media. (E) RiboSeq coverage of a conserved smORF (chr6:148354710-148355199) identified in both primary differentiated brown and subcutaneous white adipocytes (F) DIA-MS quantification of microproteins in the whole-cell lysates and secretomes of both the differentiated brown and subcutaneous white adipocyte cultures showing 4 microproteins quantified in both the whole cell lysates and secretomes that includes microproteins from the smORFs: chr6:148354710-148355199 (seen in panel E RiboSeq); chr1:156615947-156616057; chr2:129306194-129306639; and chr2:6872358-6872429. (scale bar = 25 µm in Panel A, statistics performed with paired t-test where * = p-value < 0.05, ** = p-value < 0.01, and *** = p-value < 0.001).

Having established the robustness of the data with known genes, we turned our attention to the microproteins from within these cultures. First, we fractionated the whole cell lysates and the secretome proteins as depicted in Fig. 2. We identified a total of 55 microproteins from a combined database search of the fractionated cell proteomes and secretomes using non-quantitative DDA analysis with a 1% peptide-level false discovery rate (Table S4 and Fig. 2). Considering the cell proteomes and conditioned media secretomes from both brown and subcutaneous white adipocytes, we quantified 12 microproteins specific to the cell proteomes, 7 microproteins specific to the secretomes, and 7 microproteins that were found in both compartments (Fig. 3D). None of these microproteins are listed in the UniProt database, and the peptides we detected for these microproteins do not map to any other sites in the proteome.

Some of the DIA-MS quantifiable microproteins showed different levels between brown and white adipocytes, either in the proteome or secretome fraction or both, similar to annotated proteins dynamics. The quantification via DIA-MS of the seven microproteins in unfractionated tryptic digests from both the proteomes and the secretomes (Fig. 3D) indicates that they are abundant. Fig. 3E shows the Ribo-Seq coverage for one of these microproteins encoded on the RPS4L pseudogene (RPS4L-MP). Fig. 3F shows the levels of RPS4L-MP in adipose whole cell lysate and secretome as well as quantification for a microprotein encoded within an annotated intronic region of v-abl Abelson murine leukemia viral oncogene 2 (“(intronic) ABL2-MP”), a microprotein from a smORF encoded on the putative lncRNA Gm14023 (“Gm14023-MP”), and a microprotein from a CUGBP Elav-like family member 2 uORF, (“(uORF) CELF2-MP”). RPS4L-MP and Gm14023-MP are both statistically unchanged between the brown and white cultures while the (intronic) ABL2-MP and the (uORF) CELF2-MP are significantly higher in the proteomes of brown adipocytes. Consistent with the observation that ATG start codons lead to higher microprotein levels, all of these microproteins are predicted to begin with methionine.

### Microproteins smORFs are transcriptionally regulated during diet-induced obesity

Having uncovered thousands of candidate translated smORFs in primary differentiated adipocytes, we next sought to identify which encoded microproteins are potentially functional in metabolism and obesity. To tackle this problem, we searched for evidence of smORF regulation during broad multi-organ metabolic changes triggered by diet-induced obesity (DIO). RNA-seq data was collected from BAT, epididymal white adipose tissue (eWAT), liver, scWAT, retroperitoneal fat (“Retro Fat”), and mesenteric fat (“Mesen Fat”) from cohorts of 27 week old male diet-induced obese (DIO) mice fed with high-fat diet (“HFD” for 21 weeks) and healthy age-matched control mice fed normal chow diet. To confirm that DIO mice exhibited altered metabolism, we first analyzed each tissue for differential RNA expression of annotated genes. In each of the fat depots, more than 1,000 genes showed significant changes in RNA expression in obese mice relative to healthy controls, including over 4,000 genes in eWAT (Table S5). In liver tissue, however, only 418 genes were significantly regulated. In BAT and eWAT, GO analysis revealed enrichment of lipid metabolic processes among downregulated genes and enrichment of immune system and inflammatory response pathways among upregulated genes (Fig. S2). These changes are consistent with the obesity-induced altered metabolism.

We next analyzed protein-coding smORFs expressed in adipocytes for transcriptional changes in DIO versus lean mice. In each fat tissue type, hundreds of smORFs were found to be significantly changed, while only 85 were found to be regulated in liver tissue (Fig. 4A and Fig. S3), consistent with fewer annotated genes being differentially expressed in liver relative to the fat depots. Observation of the RNA-Seq tracks for *Irs1* and *Ucp2* shows the marked differences in RNA counts over two associated smORFs (Fig. 4B) opening the possibility that these and other unannotated microproteins could function in lipid metabolism and adipocyte biology. Additionally, our Ribo-Seq data indicated ribosomal association with five predicted microproteins that are encoded on the putative lncRNA gene Brown adipose tissue enriched long non-coding RNA 1 (*Lncbate1*), which show decreased expression in several fat depots of obese mice (Fig. 4C and Table S6) (Alvarez-Dominguez et al., 2015). *Lncbate1* has been shown to be required for development and maintenance of brown adipocytes (Alvarez-Dominguez et al., 2015). Therefore, these data now call into question whether this gene’s functions are actually driven by the encoded microproteins instead of, or in conjunction with, the transcribed RNA. Lastly, a global analysis via principal component analysis (PCA) of RNA expression for smORFs from both non-uORF smORFs (Fig. 4D) as well as from uORF smORFs (Fig. S4) show that adipocyte smORFs are capable of separating out brown adipose tissue from non-BAT fat depots and liver, suggesting that also the BAT smORF expression landscape and potentially some of the microprotein levels in these tissue depots are different.

**Fig. 4.**
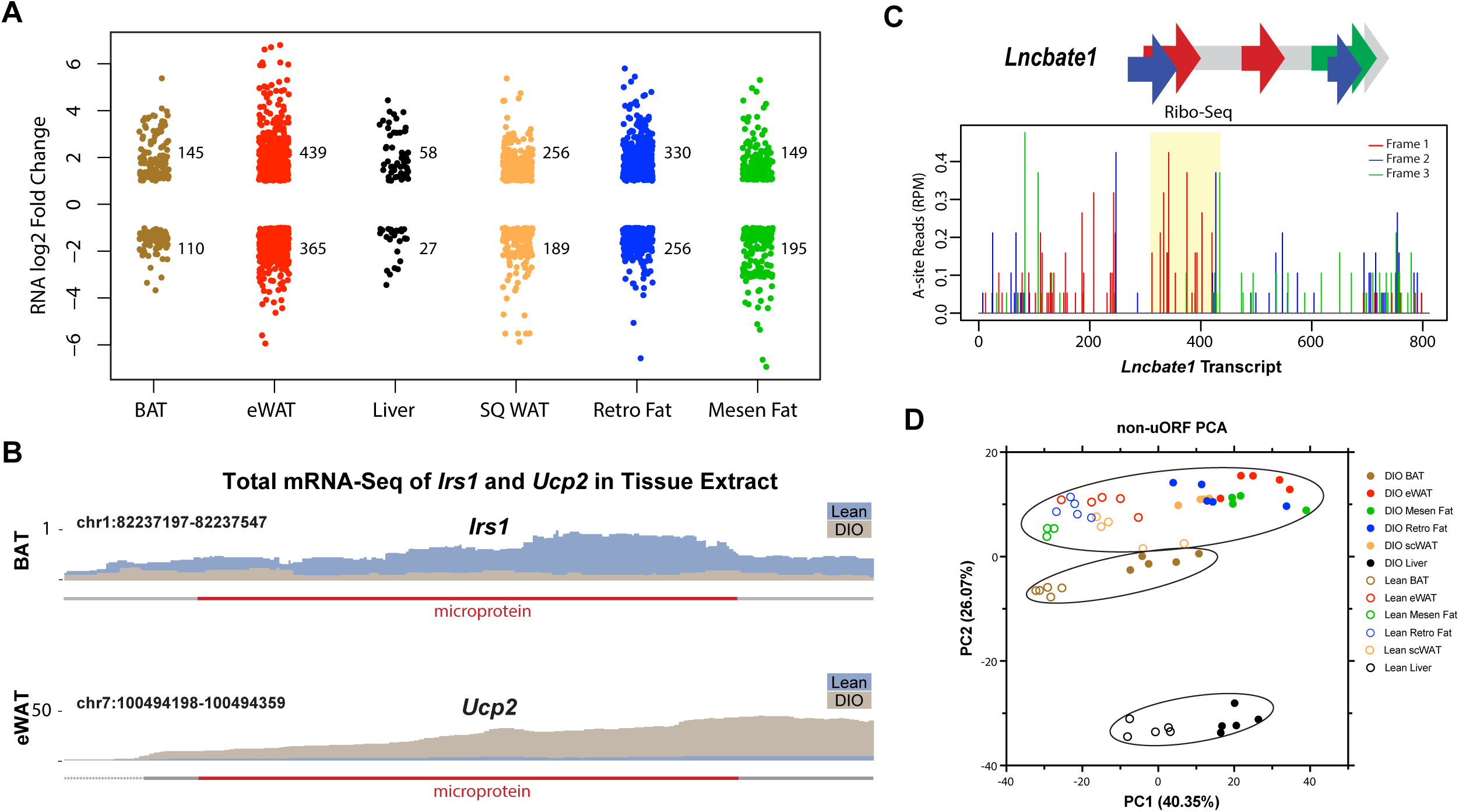
Adipose protein-coding smORFs are differentially transcribed in diet-induced obesity mice. (A) Changes in RNA expression for adipose protein-coding smORFs induced by DIO in various tissues (padj < 0.05 and |log2 fold change| ≥ 1). (B) Total mRNA-Seq read maps (equivalently scaled) in lean (blue) and DIO (brown) over the microprotein encoding smORFs of *Irs1* (comparison depicted in BAT, top) and *Ucp2* (comparison depicted in eWAT, bottom). (C) Schematic showing the locations of translated smORFs identified on *Lncbate1*. Five smORFs were identified across the primary differentiated subcutaneous white, brown, and beige adipocytes: two in frame 1, two in frame 2, and one in frame 3 (left). Representative ribosomal A-site plots (Ribo-Seq) from brown adipocytes for *Lncbate1* with one smORF identified as translated in both brown and subcutaneous white adipocytes highlighted in yellow (right). (D) PCA analysis of non-uORF protein-coding smORF RNA expression levels in tissues derived from DIO and lean mice.

### Relative quantification of age- and obesity-associated mouse plasma canonical ORF proteins and microproteins

The identification of a microprotein in the circulation is interesting as it suggests a potential novel system-wide biological functional in regulation of distant target cells or tissues. To complement our tissue RNA-Seq of obesity in mice, we analyzed plasma proteins, including microproteins, from DIO and lean mice at 26- and 41-weeks of age (sample groups shown in Fig. 5A). We used a deep fractionation DDA method to improve detection of small proteins in plasma followed by DIA-MS to quantify circulating microproteins. For the deep protein fractionation that improved our small protein coverage, we enriched the undigested plasma proteins with both C8 and C18 to enrich for the small proteome (Fig. 5B). These enriched samples were then digested with trypsin and the peptides fractionated with high-pH reverse phase (RP) fractionation (Fig. 2). The results of all of the DDA-based plasma fractionation provided evidence for 33 circulating microproteins that are each represented in the library by a single tryptic peptide (Table S7) highlighting the need for DIA-MS quantification that is robust on the single peptide level proteome-wide.

**Fig. 5.**
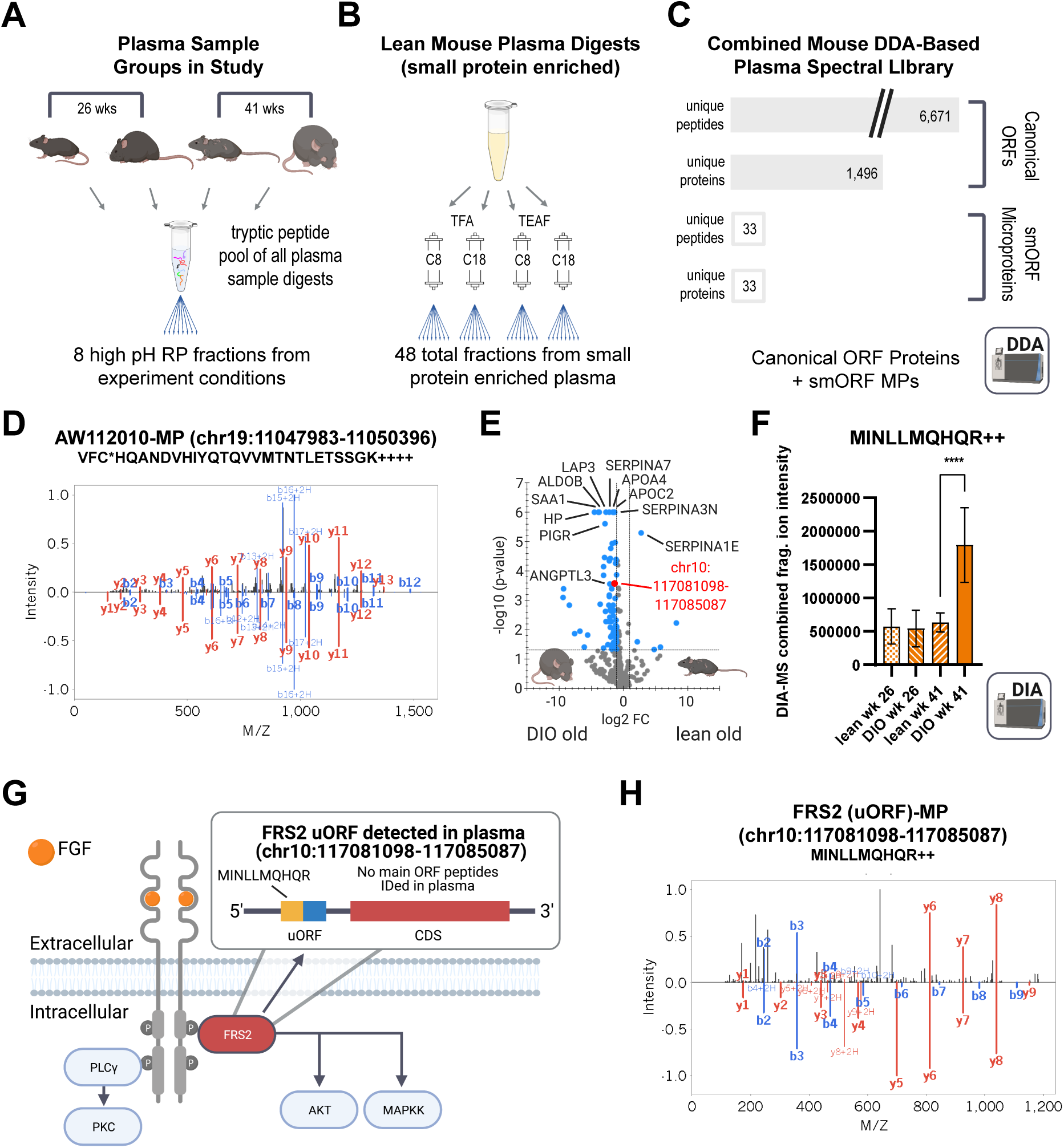
Plasma proteomics of aged obese mice identifies evidence for multiple smORF-encoded microproteins with one FRS2 uORF circulating at a higher level in the aged obese state. (A) Experimental design for plasma proteomics of 26-week and 41-week old mice with both DIO and lean groups in both ages (n=12 per group). (B) Strategy for non-quantitative DDA-based deep identification of canonical ORF proteins and smORF-encoded microproteins in lean mouse plasma shows a combinatorial fractionation strategy designed to enrich small proteins using both C8 and C18 based fractionation with both trifluoroacetic acid (TFA) and triethylammonium formate (TEAF) ion-pairing agents. Additionally, plasma from the experiment depicted in Panel A was fractionated (not pictured) with high pH RP fractionation as depicted in Figure 2. (C) Summary of all proteins identified with the fractionation strategies outlined in Panel B showing peptides and proteins identified from both the canonical ORFs and smORF-encoded microproteins. (D) Annotated MS2 fragment ion spectrum of a peptide (VFC*HQANDVHIYQTQVVMTNTLETSSGK++++, *=reduced and alkylated cysteine) that maps to a microprotein generated from the lincRNA AW112010 (chr19:11047983-11050396) discovered in the circulation via the plasma fractionation depicted in Panel B. The peptide is depicted in a butterfly plot with the measured MS2 spectrum above the x-axis and the Prosit-predicted fragment ion pattern below the x-axis (E) Volcano plot of DIA-MS quantification of the experiments depicted in Panel A comparing the DIO old condition to the lean old condition depicting regulated canonical ORF proteins along with a regulated smORF-encoded microprotein that correlates to chr10:117081098-117085087. (F) Quantification with DIA-MS of a tryptic peptide (sequence: MINLLMQHQR++) for the smORF-encoded microprotein from chr10:117081098-117085087 across all of the biological conditions in Panel A. (G) The amino acid sequence of the smORF-encoded microprotein from chr10:117081098-117085087 that maps to the uORF region of fibroblast receptor substrate 2 (FRS2) with the identified tryptic peptide (sequence: MINLLMQHQR) depicted in yellow and the whole smORF in blue/yellow. (H) Annotated MS2 fragment ion spectrum of the tryptic peptide from the FRS2 uORF (sequence: MINLLMQHQR++) was used to quantify this microprotein in Panels E and F with DIA-MS. (statistics performed with one-way ANOVA where **** = p-value < 0.0001).

Of the DDA-MS-identified microproteins, three had a predicted signal peptide that scored above the positive thresholds (>0.4 in SignalP 5.0) (Fig. S5), highlighting the ability of this workflow to identify novel secreted microproteins, but also suggesting other secretion mechanisms are at play. One of microproteins with a high SignalP score is from a smORF on a non-coding RNA (symbol A530053G22Rik) and a second is on a pseudogene described as lymphocyte antigen 6 complex pseudogene (symbol 9030619P08Rik). The presence of the latter gene provided evidence that this pseudogene is translated into a stable, circulating microprotein. This microprotein shares 50-60% sequence identity to several other mouse lymphocyte antigen 6 complex members, a family with emerging roles in immunity and cancer (Upadhyay, 2019). The third detected secreted microprotein with a confidently predicted signal peptide, AW112010-MP (Fig. 5D), was recently characterized with a role in mucosal immunity (Jackson et al., 2018; Yang et al., 2020). Our data provides new compelling evidence that AW112010-MP is indeed circulating beyond the local environment of the gut and inflammatory macrophages where it was discovered and suggests potential additional roles for this microprotein in the inflammation of obesity. Broadly, the finding of immune related genes is consistent with models of obesity and diabetes leading to inflammation and changes in immunity.

Quantitative DIA-MS comparison of different combinations of lean, obese, young, and old mice identified many canonical ORF proteins that are changed in these different physiological settings. First, we validated our quantitation with known human biomarkers of obesity-related liver disease (ALDOB, LAP3, SAA1, and PIGR) (Niu et al., 2019) and aging (HP, APOC2, APOA4, SERPINA7, and SERPINA3N) (Fig. 5E). Taken together, this data strongly shows an aged, obese mouse with numerous markers of liver injury and dysfunction (Niu et al., 2019). Alongside these established markers, we find increased levels of a 24-amino acid microprotein encoded by a smORF in the 5’UTR of fibroblast growth factor receptor substrate 2 (uORF) FRS2-MP in aged/obese mice (Fig. 5F-H). FRS2 is an intracellular protein that links ligand-activated fibroblast growth factor receptor 1 (FGFR1) to downstream signal transduction pathways (Rabin et al., 1993; Xu et al., 1998) (Fig. 5G). We did not detect evidence for the canonical FRS2 protein in the plasma. The tryptic peptide for this (uORF) FRS2-MP was MINLLMQHQR++ with its well annotated DDA-based spectrum shown in Fig. 5H. Our previous efforts in human cells identified a uORF on *FRS2* that is conserved in primates. While this MP sequence differs substantially from the mouse uORF, the presence of a uORF in both genes suggests that *FRS2* is post-transcriptionally regulated by a uORF, a hypothesis that should be investigated further.

### Ribo-Seq evidence for the translation of the predicted gene Gm8773 that encodes the mouse orexigenic FAM237B neurosecretory protein

In our Ribo-Seq data, we identified a translated smORF from the predicted gene Gm8773 (labelled as a non-coding RNA in mouse) in primary differentiated BAT- and scWAT-derived adipocyte cultures (Fig. 6A), providing the first evidence for translation of this mRNA in mice. We became interested in the Gm8773 because this 132 amino acid microprotein contains a signal peptide which would generate a 108 amino acid secreted microprotein in its final predicted processed form. Furthermore, Gm8773 is the evolutionarily conserved homolog of the human, rat, and chicken FAM237B (Ukena, 2021) which are biologically active. hFAM237B has activity against the feeding receptor GPR83, and chicken FAM237B is an orexigenic peptide that regulates feeding and adipose tissue in chickens (Ukena, 2021). The evidence for Gm8773 translation validates the translation of mFAM237B in mouse adipose cells.

**Fig. 6.**
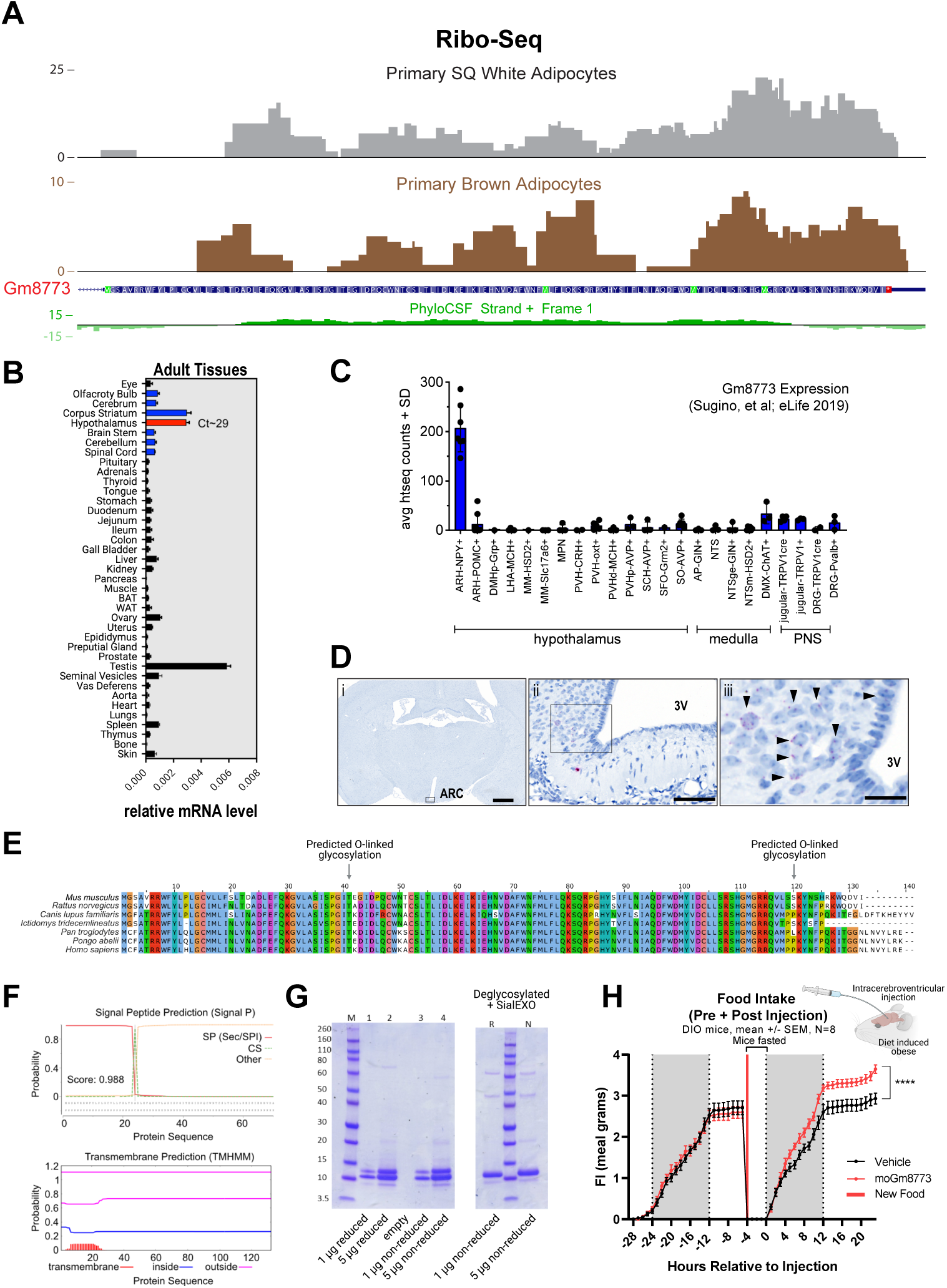
Translation, expression, and activity of Gm8773. (A) RiboSeq evidence for the translation of Gm8773 in both differentiated brown and subcutaneous white adipocytes. (B) Relative comparative tissue level mRNA expression with qPCR of Gm8773 across a panel of mouse tissues (C) Replotting of published transcriptional co-expression data of Gm8773 expression within specific nuclei of the hypothalamus and other regions of the brain; Note Gm8773 co-expressed with NPY containing neurons (Sugino et al., 2019) (D) *in situ* hybridization of Gm8773 mRNA localization to the arcuate nucleus of the hypothalamus, showing section morphometry (D.i), the arcuate nucleus and median eminence (D.ii), and arrows denoting Mouse-Gm8773+ cells in the mediobasal hypothalamus. Scale bars = 500 μm (D.i), 250 μm (D.ii), 50 μm (D.iii). (E) Amino acid level conservation of the predicted protein sequence of Gm8773 along with the two residues that have a positive O-linked glycosylation prediction score (NetOGlyc 4.0) (F) Signal peptide (Signal P 5.0) and transmembrane domain (TMHMM 2.0) predictions for Gm8773 predicted protein sequence. (G) Expression of recombinant Gm8773 in HEK cells showing a double band at the molecular weight of a protein monomer in both reducing and non-reducing conditions (left side) along with the collapsing of the Gm8773 doublet protein band down to a single band upon treatment with an O-linked glycosylation deglycosylating enzyme (right side). (H) Increased food intake is observed in mice following the intracerebroventricular (ICV injection) administration of the recombinant Gm8773 protein from Panel G (statistics performed with two-way ANOVA with significant time by treatment interaction where **** = p-value < 0.0001).

Expression and regulation provides further evidence that Gm8773 may be active in regulating mouse feeding. Previously, transcriptional control of Gm8773 was seen during single-cell RNA-Seq (scRNA-Seq) measurements of the arcuate nucleus of the hypothalamus where it is co-expressed with agouti-related peptide (AgRP) (Campbell et al., 2017). In that scRNA-Seq atlas of the arcuate nucleus, the AgRP-Gm8773 cluster is distinct from the AgRP-somatostatin cluster suggesting its potential role in somatostatin-independent AgRP-related feeding mechanisms. We further identified the transcript as tissue-specifically elevated in the hypothalamus in qPCR quantitation of the mRNA across a panel of mouse tissues (Fig. 6B). Additionally, reanalysis of published flow cytometry data of sorted cell types explanted from genetically engineered mouse strains with GFP-tagged brain loci-specific reporter genes shows clear co-expression of Gm8773 with the NPY labeled cells (Fig. 6C) (Sugino et al., 2019). These data are consistent with our morphological distribution of Gm8773 that we observe by *in situ* hybridization in hypothalamic brain slices (Fig. 6D). Together, the data strongly support an association between Gm8773 with the neuropeptide Y (NPY) / AgRP feeding locus in the arcuate nucleus, consistent with the homology and activity to FAM237B.

We reasoned that the combined evidence of the Ribo-Seq-derived protein coding potential of Gm8773/FAM237B mixed with the arcuate nucleus association of the transcript created a logical experimental hypothesis for testing a recombinantly expressed Gm8773/FAM237B protein directly in the hypothalamus. Sequence analysis predicts two O-linked glycosylation sites (Fig. 6E) (Hansen et al., 1998), no transmembrane domains (Fig. 6F) (Sonnhammer et al., 1998), and the presence of a signal peptide (Fig. 6F) (Almagro Armenteros et al., 2019) all of which support the secretion of this microprotein from cells. Recombinant expression of Gm8773/FAM237B in mammalian cells (HEK 293T) generated a protein of the correct molecular weight that ran as a monomer in SDS-PAGE despite the presence of a free cysteine (Fig 6G). Treatment of the recombinant protein with an O-linked specific deglycosylation enzyme resulted in the collapse of two proximal bands into a single band, consistent with the predicted glycosylation of Gm8773/FAM237B (Fig. 6G). The straightforward production of the recombinant Gm8773/FAM237B microprotein highlights a single example of potentially many unannotated, stable, secreted, post-translationally modified, and folded microproteins in our adipose smORF datasets.

To test the hypothesis that the mFAM237b microprotein modulates food intake, we injected our recombinant microprotein into DIO mice via intracerebroventricular (ICV) injection 30 minutes before the nocturnal cycle. We observed an induction of feeding in the mFAM237B-treated mice (Fig. 6H), consistent with expression in the feeding center (Campbell et al., 2017; Sugino et al., 2019) and homology to chicken FAM237B (Ukena, 2021), which showed similar orexigenic activity in chickens. Thus, the Ribo-Seq data validated the translation of Gm8773 to produce the mouse FAM237B homolog, which has orexigenic activity and establishes this molecule as potentially part of the brain-adipose axis to control feeding. The advantage of having established this system in mice is that we can use the large repository of knockout mice and engineering tools in mouse neuroscience and physiology to better study the role of this microprotein than would have been feasible in chickens or other organisms.

## DISCUSSION

The growing global health burden of obesity is of major concern for the future of humans. Obese individuals are at increased risk for type 2 diabetes (T2D) as well as obesity-related liver diseases such as non-alcoholic fatty liver disease (NAFLD) and non-alcoholic steatohepatitis (NASH) where cirrhosis can create the need for liver transplant in the latter and where therapeutic strategies are currently lacking (Cassidy and Syed, 2016; Newsome et al., 2021; Vuppalanchi et al., 2021). Additionally, obesity is a well-established driving risk factor of cardiometabolic health problems and cardiac-related death and has a high association with numerous cancers potentially caused by the altered systemic endocrine function created by the obese state. Nevertheless, gaps remain in our knowledge about the molecular determinants of obesity and obesity-related diseases.

While the aim of whole genome mapping projects has been to identify and validate all possible protein-coding genes, the last decade has revealed a blind spot for smORFs and microproteins, with thousands of these peptides and small proteins being recently discovered. Since we cannot study what we do not know, the discovery of smORFs and their correlate microproteins represents an important advancement in the understanding of the genome and proteome. Most microprotein annotation has focused primarily on cell lines and primary cells in culture, but an excellent example of tissue-based smORF discovery in the human heart was recently published (van Heesch et al., 2019) unlocking new smORF genes with potential function in cardiac biology and disease. With any of these newly identified smORF genes, it is necessary to perform ribosome profiling on cells or tissues of interest because the ability of these genes to be transcribed or translated in all tissues is not known.

The mouse is an ideal model for metabolism research because countless results from mouse studies have translated into humans, and the prevalence of different genetic models facilitates mechanistic studies. Adipose depots have well-documented endocrine functions with leptin, a potent circulating metabolic factor adipokine, being a long-standing exemplar (Ahima and Flier, 2000), which piqued our interest in the possibility of identifying novel secreted microproteins in adipocytes. While tissue ribosome profiling is feasible, low overall yields of RNA in adipocytes creates challenges in tissue-based Ribo-Seq and profiling endogenous adipose tissue would result in a mixture of cell types. For these reasons, we focused our Ribo-Seq efforts on primary white, beige, and brown adipocytes. This led to the identification of 3,877 smORFs that all potentially encode microproteins that are absent from the SwissProt UniProt reviewed proteome database, providing an entirely new set of adipose proteins to study and understand. We imagine that these genes can and should be included in large-scale genomic screening efforts like using CRISPR to identify novel functions as transcriptional regulators (Dumesic et al., 2019) and/or functional microproteins (Chen et al., 2020; Chu et al., 2019).

To understand the potential impact of large groups of microproteins on metabolism, we examined smORF regulation and found hundreds that are transcriptionally regulated in various adipose tissue depots in mice fed high fat diet. These putative microproteins along with the conserved smORFs represent a subset that should be at the top of any list for subsequent functional studies. More generally, with thousands of available RNA-Seq datasets, researchers can map these smORFs to different studies to identify smORFs regulated under different metabolic conditions or genetic models where changes in microprotein expression levels can lead to new testable hypotheses.

In general, microproteins are derived from mRNAs or mRNA regions thought to be non-coding—e.g., 5’- and 3’-UTRs and non-coding RNAs. With mRNAs with multipole ORFs, the smORFs are thought to act as translational regulators of the main ORFs (Chen et al., 2010; Lee et al., 2009; Wu et al., 2020). The uORF regulator of PGC1*α* is one example of an important translational regulator where the uORF generates a stable microprotein that in part controls translation of the canonical ORF (Dumesic et al., 2019) and a recent study showed a translated uORF microprotein that can inhibit PKC phosphorylation (Jayaram et al., 2021). Of course, many of these smORFs might regulate translation via ribosome stalling or some other untranslated mechanism *and* also produce a functional microprotein (Chen et al., 2020). Therefore, it is interesting to find smORFs on the mRNAs of several metabolic genes (e.g., INSR, PLIN1, UCP1), as these underappreciated genetic elements might have a key role in the positive or negative regulation of the translation of these genes, or indeed produce bioactive microproteins. Either way, these findings lead to new hypotheses and open up new avenues of inquiry into these genes. Similarly, the discovery that some functional “non-coding” RNAs, such as LNCBATE1 may in fact be translated (Alvarez-Dominguez et al., 2015), may lead to the expansion of our understanding of their biology as potential protein-coding genes and not just non-coding RNAs although the possibility of both biological functions co-occurring also exists. For metabolism, certainly the Ribo-Seq-based discovery of translation of a non-coding RNA whose amino acid sequence has a predictable secretion signal peptide, is of high interest for follow up studies such as those shown here with the observed central feeding control associated with our recombinant Gm8773.

As new genes, antibodies to microproteins need to be developed, which can be difficult and time consuming (Chu et al., 2019). Proteomics offers a solution to microprotein detection and complements Ribo-Seq by reporting on the stability and location of a microprotein. However, owing to their length, microproteins produce few tryptic peptides for detection, and the tryptic peptides they generate are often not unique to the microprotein, or none of the tryptic peptides are detectable because of their biophysical properties. These limitations hamper the validation of microproteins and have also limited previous proteomics-based quantification efforts. DIA-MS provides a more reliable detection and quantitation platform for dealing with single peptides by mass spectrometry, resulting in an ideal strategy for microproteins. Indeed, using DIA-MS, we were able to detect and quantify microproteins in our primary adipocyte cultures. Also, with the ability of proteomics to inform on the location of a microprotein, we revealed the existence of several circulating microproteins in mouse plasma, including a well-annotated spectrum for AW112010-MP, a recently discovered mouse microprotein with a role in gut immunity. By revealing AW112010-MP as a circulating microprotein it suggests a broader role for this microprotein in immunity that compliments the original discovery of the protein (Jackson et al., 2018).

At the outset of these studies Gm8773 was listed as a non-coding RNA and ribosome profiling provided the first empirical evidence that Gm8773 produces a protein in mice (this misannotation was recently revised and Gm8773 is currently listed as a protein-coding gene in RefSeq, but a “Predicted gene” in UniProt). As a result of its misannotation, there are no reports of Gm8773 translation or function, but, fortunately, homology to a characterized chicken protein FAM237B led us to show that this gene makes a stable, folded, secretable, and glycosylated recombinant microprotein that regulates food intake when administered centrally to DIO mice, which is also consistent with published chicken and rat experiments (Ukena, 2021). To compliment the central food intake increase effects seen here, the Gm8773 gene was previously reported to be co-expressed with AgRP, a known orexigenic peptide, in the hypothalamus in a scRNA-Seq atlasing effort (Campbell et al., 2017). Given the availability of readily available genetic mouse models and experimental paradigms in mice, research into the potential impact of this gene can now accelerate with more confidence aided by the verification of its protein function in mice. More generally, this example highlights the value of microprotein annotation to uncover these novel proteins and facilitate their characterization to provide new biological insights and therapeutic opportunities.

Altogether, the smORF database generated in this study will find value in mining other -omics datasets including: genomics data, bulk RNA-Seq data from disease models, proteomics data by amending protein databases with microprotein sequences, and the ever-expanding single cell datasets. Moreover, with DIA-MS adoption increasing in proteomics due to superior quantitation, we imagine that the integration of sequences will reveal new microproteins that are increased or decreased under different conditions using DIA-MS. These -omics studies will lead to targeted hypothesis driven experiments to look at the function of specific smORFs or microproteins based on their location or regulation. For instance, upstream and downstream smORFs on genes such as *Ucp1* and *Irs1* suggest a layer of translational regulation that is currently not well understood. Thus, the impact of this work will be the identification of new genes with roles in murine metabolism, and eventually some of these genes will provide insights that can provide therapeutic opportunities in obesity and diabetes and potentially other diseases.

## Supporting information

All Supplemental text and figures

## LIMITATIONS OF THIS STUDY

In this study, we provide evidence for the translation of thousands of smORFs in primary brown and white adipocytes, and the proteomic detection of microproteins from cell proteomes, conditioned media, and plasma. Mass spectrometry underestimates microprotein numbers because these polypeptides are short and generate few detectable tryptic peptides. Furthermore, without trypsin microproteins do not ionize well, which partially explains the difference in numbers between mass spectrometry and ribosome profiling. That said, smORF annotation by ribosome profiling does not guarantee that the encoded microproteins are stable, long-lived molecules capable of regulating biology, though our hypothesis is that there are enough biologically active microproteins within this pool to make it worthwhile to interrogate these genes. Indeed, the ribosome profiling data provided the first empirical evidence for the translation of the biologically active Gm8773 microprotein. Pharmacological experiments with the recombinant Gm8773 microprotein and ICV injections reveal an intriguing feeding phenotype. To determine whether these pharmacological experiments define a physiological role of Gm8773, subsequent studies should employ mouse genetics that knockout/knockin Gm8773 in cells and tissues of a living animal.

## AUTHOR CONTRIBUTIONS

C.A.B., A.J.W., M.J.M. and A.S. (Saghatelian). study conceptualization. C.A.B., A.J.W., A.L.B., S.A.M., D.F., Y.Z., and K.B. conceptualized the animal studies. T.F.M., M.N.S., C.D., J.M.V., Y.Z. A.L.B., and C.L. performed the RNA experiments and/or Ribo-Seq analysis methodologies. T.F.M., M.N.S., A.L.B., S.A.M., and C.A.B. performed the bioinformatics methodologies, visualization, and data curation work. C.A.B. and B.K. performed the mass spectrometry. C.A.B., L.K.P., B.C.S., and M.J.M. performed the mass spectrometry bioinformatics methodologies, visualization, and data curation. S.L-A., A.A., T.K., A.M., and A.S. (Simon) performed the protein production methodologies. A.J.M. performed the *in situ* hybridization methodologies. A.B. (Baquero), K.B., D.F., J.H., R.D., S.P., C.S., and G.Z. performed the *in vivo* animal studies methodologies. C.A.B., M.J.M., and A. S. (Saghatelian) performed supervision of the research. T.F.M., C.A.B., and A. S. (Saghatelian) performed writing of the original draft.

## ACKNOWLEDGEMENTS

The authors would like to thank Erin Whalen, Fiona McMurray, and Nick Cox for continuous guidance, support, enthusiasm, and critical feedback throughout the planning and writing of this work. Additionally, the authors would like to thank Kevin Grove and Mads Tang-Christensen for support for the program. The laboratory of Michael J. MacCoss would like to acknowledge financial support in part from the National Institutes of Health grants P41 GM103533, R24 GM141156, and U19 AG065156 and has additional support from a sponsored research agreement with Novo Nordisk Research Center Seattle, Inc. The laboratory of Alan Saghatelian would like to acknowledge financial support in part from the NIH P30CA014195, R01GM102491, and RC2DK129961 and also has additional support from a sponsored research agreement with Novo Nordisk Research Center Seattle, Inc.

## SUPPLEMENTARY MATERIALS

Materials and Methods

Key Resources Table

Supplementary Figures S1-S5

Supplementary Tables S1-S7

## REFERENCES

Ahima, R.S., and Flier, J.S. (2000). Leptin. Annu. Rev. Physiol. 62, 413–437.

Almagro Armenteros, J.J., Tsirigos, K.D., Sønderby, C.K., Petersen, T.N., Winther, O., Brunak, S., von Heijne, G., and Nielsen, H. (2019). SignalP 5.0 improves signal peptide predictions using deep neural networks. Nat. Biotechnol. 37, 420–423.

Alvarez-Dominguez, J.R., Bai, Z., Xu, D., Yuan, B., Lo, K.A., Yoon, M.J., Lim, Y.C., Knoll, M., Slavov, N., Chen, S., et al. (2015). De Novo Reconstruction of Adipose Tissue Transcriptomes Reveals Long Non-coding RNA Regulators of Brown Adipocyte Development. Cell Metab 21, 764–776.

Amodei, D., Egertson, J., MacLean, B.X., Johnson, R., Merrihew, G.E., Keller, A., Marsh, D., Vitek, O., Mallick, P., and MacCoss, M.J. (2019). Improving Precursor Selectivity in Data-Independent Acquisition Using Overlapping Windows. J. Am. Soc. Mass Spectrom. 30, 669–684.

Anderson, D.M., Anderson, K.M., Chang, C.L., Makarewich, C.A., Nelson, B.R., McAnally, J.R., Kasaragod, P., Shelton, J.M., Liou, J., Bassel-Duby, R., et al. (2015). A micropeptide encoded by a putative long noncoding RNA regulates muscle performance. Cell 160, 595–606.

Andreadis, A., Hsu, Y.P., Kohlhaw, G.B., and Schimmel, P. (1982). Nucleotide sequence of yeast LEU2 shows 5’-noncoding region has sequences cognate to leucine. Cell 31, 319–325.

Ashburner, M., Ball, C.A., Blake, J.A., Botstein, D., Butler, H., Cherry, J.M., Davis, A.P., Dolinski, K., Dwight, S.S., Eppig, J.T., et al. (2000). Gene ontology: tool for the unification of biology. The Gene Ontology Consortium. Nat Genet 25, 25–29.

Bairoch, A., and Apweiler, R. (1997). The SWISS-PROT protein sequence data bank and its supplement TrEMBL. Nucleic Acids Res. 25, 31–36.

Bartelt, A., John, C., Schaltenberg, N., Berbée, J.F.P., Worthmann, A., Cherradi, M.L., Schlein, C., Piepenburg, J., Boon, M.R., Rinninger, F., et al. (2017). Thermogenic adipocytes promote HDL turnover and reverse cholesterol transport. Nat. Commun. 8, 15010.

Bi, P., Ramirez-Martinez, A., Li, H., Cannavino, J., McAnally, J.R., Shelton, J.M., Sanchez-Ortiz, E., Bassel-Duby, R., and Olson, E.N. (2017). Control of muscle formation by the fusogenic micropeptide myomixer. Science 356, 323–327.

Bookout, A., and Mangelsdorf, D. (2003). Quantitative real-time PCR protocol for analysis of nuclear receptor signaling pathways. Nucl. Recept. Signal.

Bookout, A.L., Jeong, Y., Downes, M., Yu, R.T., Evans, R.M., and Mangelsdorf, D.J. (2006a). Anatomical Profiling of Nuclear Receptor Expression Reveals a Hierarchical Transcriptional Network. Cell 126, 789–799.

Bookout, A.L., Cummins, C.L., Mangelsdorf, D.J., Pesola, J.M., and Kramer, M.F. (2006b). High-Throughput Real-Time Quantitative Reverse Transcription PCR. Curr. Protoc. Mol. Biol. 73, 15.8.1-15.8.28.

Calvo, S.E., Pagliarini, D.J., and Mootha, V.K. (2009). Upstream open reading frames cause widespread reduction of protein expression and are polymorphic among humans. Proc Natl Acad Sci U A 106, 7507–7512.

Campbell, J.N., Macosko, E.Z., Fenselau, H., Pers, T.H., Lyubetskaya, A., Tenen, D., Goldman, M., Verstegen, A.M.J., Resch, J.M., McCarroll, S.A., et al. (2017). A molecular census of arcuate hypothalamus and median eminence cell types. Nat. Neurosci. 20, 484–496.

Cannon, B., and Nedergaard, J. (2001). Cultures of adipose precursor cells from brown adipose tissue and of clonal brown-adipocyte-like cell lines. Methods Mol Biol 155, 213– 224.

Cannon, B., and Nedergaard, J. (2004). Brown Adipose Tissue: Function and Physiological Significance. Physiol. Rev. 84, 277–359.

Cassidy, S., and Syed, B.A. (2016). Nonalcoholic steatohepatitis (NASH) drugs market. Nat. Rev. Drug Discov. 15, 745–746.

Chambers, M.C., Maclean, B., Burke, R., Amodei, D., Ruderman, D.L., Neumann, S., Gatto, L., Fischer, B., Pratt, B., Egertson, J., et al. (2012). A cross-platform toolkit for mass spectrometry and proteomics. Nat. Biotechnol. 30, 918–920.

Chen, J., Brunner, A.D., Cogan, J.Z., Nunez, J.K., Fields, A.P., Adamson, B., Itzhak, D.N., Li, J.Y., Mann, M., Leonetti, M.D., et al. (2020). Pervasive functional translation of noncanonical human open reading frames. Science 367, 1140–1146.

Chen, Y.-J., Tan, B.C.-M., Cheng, Y.-Y., Chen, J.-S., and Lee, S.-C. (2010). Differential regulation of CHOP translation by phosphorylated eIF4E under stress conditions. Nucleic Acids Res. 38, 764–777.

Chu, Q., Martinez, T.F., Novak, S.W., Donaldson, C.J., Tan, D., Vaughan, J.M., Chang, T., Diedrich, J.K., Andrade, L., Kim, A., et al. (2019). Regulation of the ER stress response by a mitochondrial microprotein. Nat Commun 10, 4883.

Chugunova, A., Loseva, E., Mazin, P., Mitina, A., Navalayeu, T., Bilan, D., Vishnyakova, P., Marey, M., Golovina, A., Serebryakova, M., et al. (2019). LINC00116 codes for a mitochondrial peptide linking respiration and lipid metabolism. Proc Natl Acad Sci U A 116, 4940–4945.

Dalbøge, L.S., Jacobsen, J.M., Mehrotra, S., Mercer, A.J., Cox, N., Liu, F., Bennett, C.M., Said, M., Tang-Christensen, M., Raun, K., et al. (2021). Evaluation of VGF peptides as potential anti-obesity candidates in pre-clinical animal models. Peptides 136, 170444.

Deutsch, E.W., Mendoza, L., Shteynberg, D., Slagel, J., Sun, Z., and Moritz, R.L. (2015). Trans-Proteomic Pipeline, a standardized data processing pipeline for large-scale reproducible proteomics informatics. Proteomics Clin. Appl. 9, 745–754.

Dobin, A., Davis, C.A., Schlesinger, F., Drenkow, J., Zaleski, C., Jha, S., Batut, P., Chaisson, M., and Gingeras, T.R. (2013). STAR: ultrafast universal RNA-seq aligner. Bioinformatics 29, 15–21.

Dumesic, P.A., Egan, D.F., Gut, P., Tran, M.T., Parisi, A., Chatterjee, N., Jedrychowski, M., Paschini, M., Kazak, L., Wilensky, S.E., et al. (2019). An Evolutionarily Conserved uORF Regulates PGC1alpha and Oxidative Metabolism in Mice, Flies, and Bluefin Tuna. Cell Metab 30, 190–200 e6.

Elias, J.E., and Gygi, S.P. (2007). Target-decoy search strategy for increased confidence in large-scale protein identifications by mass spectrometry. Nat. Methods 4, 207–214.

Eng, J.K., Jahan, T.A., and Hoopmann, M.R. (2013). Comet: An open-source MS/MS sequence database search tool. PROTEOMICS 13, 22–24.

Fernandez-Costa, C., Martinez-Bartolome, S., McClatchy, D., and Yates, J.R. (2020). Improving Proteomics Data Reproducibility with a Dual-Search Strategy. Anal Chem 92, 1697–1701.

Friesen, M., Warren, C.R., Yu, H., Toyohara, T., Ding, Q., Florido, M.H.C., Sayre, C., Pope, B.D., Goff, L.A., Rinn, J.L., et al. (2020). Mitoregulin Controls beta-Oxidation in Human and Mouse Adipocytes. Stem Cell Rep. 14, 590–602.

Fu, M., Sun, T., Bookout, A.L., Downes, M., Yu, R.T., Evans, R.M., and Mangelsdorf, D.J. (2005). A Nuclear Receptor Atlas: 3T3-L1 Adipogenesis. Mol. Endocrinol. 19, 2437–2450.

Gertz, E.M., Yu, Y.-K., Agarwala, R., Schäffer, A.A., and Altschul, S.F. (2006). Composition-based statistics and translated nucleotide searches: improving the TBLASTN module of BLAST. BMC Biol. 4, 1–14.

Gessulat, S., Schmidt, T., Zolg, D.P., Samaras, P., Schnatbaum, K., Zerweck, J., Knaute, T., Rechenberger, J., Delanghe, B., Huhmer, A., et al. (2019). Prosit: proteome-wide prediction of peptide tandem mass spectra by deep learning. Nat. Methods 16, 509–518.

Gillet, L.C., Navarro, P., Tate, S., Röst, H., Selevsek, N., Reiter, L., Bonner, R., and Aebersold, R. Targeted Data Extraction of the MS/MS Spectra Generated by Data-independent Acquisition: A New Concept for Consistent and Accurate Proteome Analysis | Elsevier Enhanced Reader. Mol. Cell. Proteomics 11.

Griffin, E., Re, A., Hamel, N., Fu, C., Bush, H., McCaffrey, T., and Asch, A.S. (2001). A link between diabetes and atherosclerosis: Glucose regulates expression of CD36 at the level of translation. Nat Med 7, 840–846.

Hansen, J.E., Lund, O., Tolstrup, N., Gooley, A.A., Williams, K.L., and Brunak, S. (1998). NetOglyc: Prediction of mucin type O-glycosylation sites based on sequence context and surface accessibility. Glycoconj. J. 15, 115–130.

Hausman, D.B., Park, H.J., and Hausman, G.J. (2008). Isolation and culture of preadipocytes from rodent white adipose tissue. Methods Mol Biol 456, 201–219.

van Heesch, S., Witte, F., Schneider-Lunitz, V., Schulz, J.F., Adami, E., Faber, A.B., Kirchner, M., Maatz, H., Blachut, S., and Sandmann, C.-L. (2019). The translational landscape of the human heart. Cell 178, 242–260.

Heinz, S., Benner, C., Spann, N., Bertolino, E., Lin, Y.C., Laslo, P., Cheng, J.X., Murre, C., Singh, H., and Glass, C.K. (2010). Simple Combinations of Lineage-Determining Transcription Factors Prime cis-Regulatory Elements Required for Macrophage and B Cell Identities. Mol. Cell 38, 576–589.

Huang, N., Li, F., Zhang, M., Zhou, H., Chen, Z., Ma, X., Yang, L., Wu, X., Zhong, J., Xiao, F., et al. (2021). An Upstream Open Reading Frame in Phosphatase and Tensin Homolog Encodes a Circuit Breaker of Lactate Metabolism. Cell Metab 33, 128–144 e9.

Hultman, K., Scarlett, J.M., Baquero, A.F., Cornea, A., Zhang, Y., Salinas, C.B.G., Brown, J., Morton, G.J., Whalen, E.J., Grove, K.L., et al. (2019). The central fibroblast growth factor receptor/beta klotho system: comprehensive mapping in mus musculus and comparisons to non-human primate and human samples using an automated in situ hybridization platform. J. Comp. Neurol. 527, 2069–2085.

Ingolia, N.T., Ghaemmaghami, S., Newman, J.R., and Weissman, J.S. (2009). Genome-wide analysis in vivo of translation with nucleotide resolution using ribosome profiling. Science 324, 218–223.

Ingolia, N.T., Lareau, L.F., and Weissman, J.S. (2011). Ribosome profiling of mouse embryonic stem cells reveals the complexity and dynamics of mammalian proteomes. Cell 147, 789–802.

Iwaki, T., Yamashita, H., and Hayawaka, T. (2001). A Color Atlas of Sectional Anatomy (Tokyo).

Jackson, R., Kroehling, L., Khitun, A., Bailis, W., Jarret, A., York, A.G., Khan, O.M., Brewer, J.R., Skadow, M.H., Duizer, C., et al. (2018). The translation of non-canonical open reading frames controls mucosal immunity. Nature 564, 434–438.

Jayaram, D.R., Frost, S., Argov, C., Liju, V.B., Anto, N.P., Muraleedharan, A., Ben-Ari, A., Sinay, R., Smoly, I., Novoplansky, O., et al. (2021). Unraveling the hidden role of a uORF-encoded peptide as a kinase inhibitor of PKCs. Proc. Natl. Acad. Sci. 118, e2018899118.

Ji, Z. (2018). RibORF: Identifying Genome-Wide Translated Open Reading Frames Using Ribosome Profiling. Curr Protoc Mol Biol 124, e67.

Ji, Z., Song, R., Regev, A., and Struhl, K. (2015). Many lncRNAs, 5’UTRs, and pseudogenes are translated and some are likely to express functional proteins. Elife 4, e08890.

Kajimura, S. (2015). Promoting brown and beige adipocyte biogenesis through the PRDM16 pathway. Int J Obes Suppl 5, S11–4.

Käll, L., Krogh, A., and Sonnhammer, E.L.L. (2004). A Combined Transmembrane Topology and Signal Peptide Prediction Method. J. Mol. Biol. 338, 1027–1036.

Käll, L., Canterbury, J.D., Weston, J., Noble, W.S., and MacCoss, M.J. (2007). Semi-supervised learning for peptide identification from shotgun proteomics datasets. Nat. Methods 4, 923–925.

Keller, A., Nesvizhskii, A.I., Kolker, E., and Aebersold, R. (2002). Empirical Statistical Model To Estimate the Accuracy of Peptide Identifications Made by MS/MS and Database Search. Anal. Chem. 74, 5383–5392.

Kozak, M. (2002). Pushing the limits of the scanning mechanism for initiation of translation. Gene 299, 1–34.

Kusminski, C.M., Bickel, P.E., and Scherer, P.E. (2016). Targeting adipose tissue in the treatment of obesity-associated diabetes. Nat Rev Drug Discov 15, 639–660.

Lee, C., Zeng, J., Drew, B.G., Sallam, T., Martin-Montalvo, A., Wan, J., Kim, S.J., Mehta, H., Hevener, A.L., de Cabo, R., et al. (2015). The mitochondrial-derived peptide MOTS-c promotes metabolic homeostasis and reduces obesity and insulin resistance. Cell Metab 21, 443–454.

Lee, D.S.M., Park, J., Kromer, A., Baras, A., Rader, D.J., Ritchie, M.D., Ghanem, L.R., and Barash, Y. (2021). Disrupting upstream translation in mRNAs is associated with human disease. Nat. Commun. 12, 1515.

Lee, Y.Y., Cevallos, R.C., and Jan, E. (2009). An upstream open reading frame regulates translation of GADD34 during cellular stresses that induce eIF2alpha phosphorylation. J Biol Chem 284, 6661–6673.

Liao, Y., Wang, J., Jaehnig, E.J., Shi, Z., and Zhang, B. (2019). WebGestalt 2019: gene set analysis toolkit with revamped UIs and APIs. Nucleic Acids Res. 47, W199–W205.

Lin, M.F., Jungreis, I., and Kellis, M. (2011). PhyloCSF: a comparative genomics method to distinguish protein coding and non-coding regions. Bioinformatics 27, i275– i282.

Lin, Y.F., Xiao, M.H., Chen, H.X., Meng, Y., Zhao, N., Yang, L., Tang, H., Wang, J.L., Liu, X., Zhu, Y., et al. (2019). A novel mitochondrial micropeptide MPM enhances mitochondrial respiratory activity and promotes myogenic differentiation. Cell Death Dis 10, 528.

Love, M.I., Huber, W., and Anders, S. (2014). Moderated estimation of fold change and dispersion for RNA-seq data with DESeq2. Genome Biol. 15, 550.

Ludwig, C., Gillet, L., Rosenberger, G., Amon, S., Collins, B.C., and Aebersold, R. (2018). Data-independent acquisition-based SWATH-MS for quantitative proteomics: a tutorial. Mol. Syst. Biol. 14, e8126.

Ma, J., Diedrich, J.K., Jungreis, I., Donaldson, C., Vaughan, J., Kellis, M., Yates, J.R., and Saghatelian, A. (2016). Improved Identification and Analysis of Small Open Reading Frame Encoded Polypeptides. Anal. Chem. 88, 3967–3975.

Ma, J., Saghatelian, A., and Shokhirev, M.N. (2018). The influence of transcript assembly on the proteogenomics discovery of microproteins. PLOS ONE 13, e0194518.

MacLean, B., Tomazela, D.M., Shulman, N., Chambers, M., Finney, G.L., Frewen, B., Kern, R., Tabb, D.L., Liebler, D.C., and MacCoss, M.J. (2010). Skyline: an open source document editor for creating and analyzing targeted proteomics experiments. Bioinformatics 26, 966–968.

Makarewich, C.A., Baskin, K.K., Munir, A.Z., Bezprozvannaya, S., Sharma, G., Khemtong, C., Shah, A.M., McAnally, J.R., Malloy, C.R., Szweda, L.I., et al. (2018). MOXI Is a Mitochondrial Micropeptide That Enhances Fatty Acid beta-Oxidation. Cell Rep 23, 3701–3709.

Martinez, T.F., Chu, Q., Donaldson, C., Tan, D., Shokhirev, M.N., and Saghatelian, A. (2020). Accurate annotation of human protein-coding small open reading frames. Nat Chem Biol 16, 458–468.

Miller, P.F., and Hinnebusch, A.G. (1990). cis-acting sequences involved in the translational control of GCN4 expression. Biochim Biophys Acta 1050, 151–154.

Navarro, P., Kuharev, J., Gillet, L.C., Bernhardt, O.M., MacLean, B., Röst, H.L., Tate, S.A., Tsou, C.-C., Reiter, L., Distler, U., et al. (2016). A multicenter study benchmarks software tools for label-free proteome quantification. Nat. Biotechnol. 34, 1130–1136.

Nelson, B.R., Makarewich, C.A., Anderson, D.M., Winders, B.R., Troupes, C.D., Wu, F., Reese, A.L., McAnally, J.R., Chen, X., Kavalali, E.T., et al. (2016). A peptide encoded by a transcript annotated as long noncoding RNA enhances SERCA activity in muscle. Science 351, 271–275.

Newsome, P.N., Buchholtz, K., Cusi, K., Linder, M., Okanoue, T., Ratziu, V., Sanyal, A.J., Sejling, A.-S., and Harrison, S.A. (2021). A Placebo-Controlled Trial of Subcutaneous Semaglutide in Nonalcoholic Steatohepatitis. N. Engl. J. Med. 384, 1113–1124.

Niu, L., Geyer, P.E., Wewer Albrechtsen, N.J., Gluud, L.L., Santos, A., Doll, S., Treit, P.V., Holst, J.J., Knop, F.K., and Vilsbøll, T. (2019). Plasma proteome profiling discovers novel proteins associated with non-alcoholic fatty liver disease. Mol. Syst. Biol. 15, e8793.

Orava, J., Nuutila, P., Lidell, M.E., Oikonen, V., Noponen, T., Viljanen, T., Scheinin, M., Taittonen, M., Niemi, T., Enerbäck, S., et al. (2011). Different Metabolic Responses of Human Brown Adipose Tissue to Activation by Cold and Insulin. Cell Metab. 14, 272– 279.

Paxinos, G., and Franklin, K. (2019). The Mouse Brain in Stereotaxic Coordinates, Compact.

Pino, L.K., Just, S.C., MacCoss, M.J., and Searle, B.C. (2020). Acquiring and Analyzing Data Independent Acquisition Proteomics Experiments without Spectrum Libraries. Mol. Cell. Proteomics 19, 1088–1103.

Prensner, J.R., Enache, O.M., Luria, V., Krug, K., Clauser, K.R., Dempster, J.M., Karger, A., Wang, L., Stumbraite, K., Wang, V.M., et al. (2021). Noncanonical open reading frames encode functional proteins essential for cancer cell survival. Nat Biotechnol 39, 697–704.

Rabin, S.J., Cleghon, V., and Kaplan, D.R. (1993). SNT, a differentiation-specific target of neurotrophic factor-induced tyrosine kinase activity in neurons and PC12 cells. Mol. Cell. Biol. 13, 2203–2213.

Reilly, S.M., and Saltiel, A.R. (2017). Adapting to obesity with adipose tissue inflammation. Nat Rev Endocrinol 13, 633–643.

Roberts, L.D., Bostrom, P., O’Sullivan, J.F., Schinzel, R.T., Lewis, G.D., Dejam, A., Lee, Y.K., Palma, M.J., Calhoun, S., Georgiadi, A., et al. (2014). beta-Aminoisobutyric acid induces browning of white fat and hepatic beta-oxidation and is inversely correlated with cardiometabolic risk factors. Cell Metab 19, 96–108.

Rose, M., and Botstein, D. (1983). Structure and function of the yeast URA3 gene. Differentially regulated expression of hybrid beta-galactosidase from overlapping coding sequences in yeast. J Mol Biol 170, 883–904.

Rosen, E.D., and Spiegelman, B.M. (2014). What We Talk About When We Talk About Fat. Cell 156, 20–44.

Searle, B.C., Pino, L.K., Egertson, J.D., Ting, Y.S., Lawrence, R.T., MacLean, B.X., Villén, J., and MacCoss, M.J. (2018). Chromatogram libraries improve peptide detection and quantification by data independent acquisition mass spectrometry. Nat. Commun. 9.

Searle, B.C., Swearingen, K.E., Barnes, C.A., Schmidt, T., Gessulat, S., Küster, B., and Wilhelm, M. (2020). Generating high quality libraries for DIA MS with empirically corrected peptide predictions. Nat. Commun. 11, 1548.

Shimizu, Y., Nikami, H., Tsukazaki, K., Machado, U.F., Yano, H., Seino, Y., and Saito, M. (1993). Increased expression of glucose transporter GLUT-4 in brown adipose tissue of fasted rats after cold exposure. Am. J. Physiol.-Endocrinol. Metab. 264, E890–E895.

Slavoff, S.A., Mitchell, A.J., Schwaid, A.G., Cabili, M.N., Ma, J., Levin, J.Z., Karger, A.D., Budnik, B.A., Rinn, J.L., and Saghatelian, A. (2013). Peptidomic discovery of short open reading frame-encoded peptides in human cells. Nat Chem Biol 9, 59–64.

Smith, U., and Kahn, B.B. (2016). Adipose tissue regulates insulin sensitivity: role of adipogenesis, de novo lipogenesis and novel lipids. J Intern Med 280, 465–475.

Sonnhammer, E.L., Von Heijne, G., and Krogh, A. (1998). A hidden Markov model for predicting transmembrane helices in protein sequences. pp. 175–182.

Stein, C.S., Jadiya, P., Zhang, X., McLendon, J.M., Abouassaly, G.M., Witmer, N.H., Anderson, E.J., Elrod, J.W., and Boudreau, R.L. (2018). Mitoregulin: A lncRNA-Encoded Microprotein that Supports Mitochondrial Supercomplexes and Respiratory Efficiency. Cell Rep 23, 3710–3720 e8.

Sugino, K., Clark, E., Schulmann, A., Shima, Y., Wang, L., Hunt, D.L., Hooks, B.M., Tränkner, D., Chandrashekar, J., Picard, S., et al. (2019). Mapping the transcriptional diversity of genetically and anatomically defined cell populations in the mouse brain. ELife 8, e38619.

The, M., MacCoss, M.J., Noble, W.S., and Käll, L. (2016). Fast and Accurate Protein False Discovery Rates on Large-Scale Proteomics Data Sets with Percolator 3.0. J. Am. Soc. Mass Spectrom. 27, 1719–1727.

The GBD 2015 Obesity Collaborators (2017). Health Effects of Overweight and Obesity in 195 Countries over 25 Years. N. Engl. J. Med. 377, 13–27.

Ting, Y.S., Egertson, J.D., Bollinger, J.G., Searle, B.C., Payne, S.H., Noble, W.S., and MacCoss, M.J. (2017). PECAN: library-free peptide detection for data-independent acquisition tandem mass spectrometry data. Nat. Methods 14, 903–908.

Trapnell, C., Williams, B.A., Pertea, G., Mortazavi, A., Kwan, G., van Baren, M.J., Salzberg, S.L., Wold, B.J., and Pachter, L. (2010). Transcript assembly and quantification by RNA-Seq reveals unannotated transcripts and isoform switching during cell differentiation. Nat. Biotechnol. 28, 511–515.

Ukena, K. (2021). Neurosecretory protein GL/neurosecretory protein GM. In Handbook of Hormones, (Elsevier), pp. 165–167.

Upadhyay, G. (2019). Emerging role of lymphocyte antigen-6 family of genes in cancer and immune cells. Front. Immunol. 10, 819.

Vijay, J., Gauthier, M.-F., Biswell, R.L., Louiselle, D.A., Johnston, J.J., Cheung, W.A., Belden, B., Pramatarova, A., Biertho, L., Gibson, M., et al. (2020). Single-cell analysis of human adipose tissue identifies depot- and disease-specific cell types. Nat. Metab. 2, 97–109.

Vuppalanchi, R., Noureddin, M., Alkhouri, N., and Sanyal, A.J. (2021). Therapeutic pipeline in nonalcoholic steatohepatitis. Nat. Rev. Gastroenterol. Hepatol. 18, 373–392.

Wingett, S.W., and Andrews, S. (2018). FastQ Screen: A tool for multi-genome mapping and quality control. F1000Research 7, 1338.

Wu, Q., Wright, M., Gogol, M.M., Bradford, W.D., Zhang, N., and Bazzini, A.A. (2020). Translation of small downstream ORFs enhances translation of canonical main open reading frames. EMBO J 39, e104763.

Xu, H., Lee, K.W., and Goldfarb, M. (1998). Novel Recognition Motif on Fibroblast Growth Factor Receptor Mediates Direct Association and Activation of SNT Adapter Proteins *. J. Biol. Chem. 273, 17987–17990.

Yang, X., Bam, M., Becker, W., Nagarkatti, P.S., and Nagarkatti, M. (2020). Long Noncoding RNA AW112010 Promotes the Differentiation of Inflammatory T Cells by Suppressing IL-10 Expression through Histone Demethylation. J. Immunol. 205, 987– 993.

Zhang, Q., Vashisht, A.A., O’Rourke, J., Corbel, S.Y., Moran, R., Romero, A., Miraglia, L., Zhang, J., Durrant, E., Schmedt, C., et al. (2017). The microprotein Minion controls cell fusion and muscle formation. Nat Commun 8, 15664.

