## Supplementary material for "Profiling Mouse Brown and White Adipocytes to Identify Metabolically Relevant Small ORFs and Functional Microproteins": All Supplemental text and figures

**SUPPLEMENTARY MATERIALS IN THIS DOCUMENT**

Materials and Methods
Key Resources Table
Supplementary Figures S1-S5
Supplementary Tables S1-S7

#### MATERIALS AND METHODS

#### EXPERIMENTAL MODEL AND SUBJECT DETAILS

All of the animal experiments with the exclusion of the tissue bank RNA extracts that were used for qPCR of Gm8773 were performed in accordance with internationally accepted principles for the use of laboratory animals and were approved by the Novo Nordisk Research Center Seattle Institutional Animal Care and Use Committee and the Novo Nordisk Ethical Review Committee. All functional animal studies were performed on male mice.

For the tissue bank RNA extracts that were used for qPCR of Gm8773, pure strain C57/Bl6J and 129×1/SvJ mice (6-week old male and female) were purchased from Jackson Laboratory (Bar Harbor, ME) and were allowed to reach 8–9 weeks of age on a 12 hr light/dark cycle and fed standard diets (Harlan Teklad #7001; Madison, WI). Females were housed together to ensure a synchronous estrus cycle, although that point was not determined prior to the harvest. All protocols were approved by the University of Texas Southwestern Medical Center Institutional Animal Care and Use Committee.

**Creation of diet-induced obese (DIO) mouse tissue atlas for RNA-Seq analysis**

30 Male, C57Bl/6J mice were obtained from The Jackson Laboratory (Stock #000664 Bar Harbor, ME, USA) and were assigned to high fat (DIO) or low fat (Control) diet groups at 6 weeks of age. The diets were 60% energy from fat (D12492i, Research Diets; N = 15) and 10% energy from fat (D12450Bi, Research Diets; N = 15). Sanitized water and the test diets were available ad libitum. Animals were transported to Novo Nordisk Research Center Seattle at 21 weeks of age and were maintained on the same test diets for an additional 6 weeks in a light and temperature-controlled room (12/12 light-dark cycle, 22 +/- 2°C). At 27 weeks of age the group mean body weights were 47.8g +/- 2.8 g (DIO) and 31.2g +/- 1.3 g (Control). Animals were euthanized via CO2 inhalation approximately 3 to 8 hours into the start of the light period; with food removal 2 hours prior to euthanasia. The order of euthanasia and tissue collection was alternated between groups to control for bias. Adipose and liver tissues (and other tissues not reported here) were removed rapidly, frozen in liquid nitrogen and stored at -80°C until processed for RNA. All animal experiments were performed in accordance with internationally accepted principles for the use of laboratory animals and were approved by the Novo Nordisk Research Center Seattle Institutional Animal Care and Use Committee and the Novo Nordisk Ethical Review Committee.

**Creation of DIO/lean young/old mouse plasma samples**

Male C57/BL6J mice were purchased from Jackson and grouped housed in a 12 h light/dark cycle with lights on at 6 AM and with *ad libitum* access to food and water throughout the experiment. The diet-induced obese (DIO) groups represented 45% of the original animals in the cohort that were kept on chow (PicoLab Rodent Diet 20) diet for their first 4 weeks after arrival and subsequently switched to high fat diet (45% HFD Research Diets: D12451i) at week 10 with these DIO animals remaining on the HFD for the duration of the experiment. The chow (PicoLab Rodent Diet 20) fed animals remained on the same diet throughout the entire experiment. Plasma was collected from both DIO and lean (chow) animal groups that were harvested at either 26- or 41-weeks of age representing the “young” and “old” groups, respectively. Each group represents 8 animals (n=8). For plasma collections, animals were euthanized with asphyxiation with $CO_2$ followed by cardiac puncture with individual blood collections into Li Heparin tubes (BD Microtainer tubes cat# 365965)). 120µl of plasma was separated for each sample by centrifugation at 10,000 RCF for 4 minutes $4^0$C followed by transferring the plasma supernatant into fresh tubes and immediately snap freezing the aliquots in liquid nitrogen.

**Intracerebroventricular (ICV) administration of Gm8773 protein product for assessing food intake and body weight effects in the diet-induced obese mouse**

ICV experiments were performed as described previously (Dalbøge et al., 2021). Briefly, a permanent 26-gauge-guided cannula (C315GS-4/SPC, PlasticsOne, Roanoke, VA, USA) projecting to the lateral ventricle was implanted into DIO mice which were male C57BL/6J mice purchased from Jackson Laboratories with obesity induced in-house via continuous feeding with high fat diet (45% HFD Research Diets: D12451i) from age 10 weeks on. DIO mice were cannulated for ICV injections at age 51 weeks. The surgeries were performed as follows: Isoflurane anesthesia was administered to Mice who were placed in a stereotaxic instrument that was maintained at 36−37 °C throughout surgery on a heating pad. Continuous isoflurane was administered using a specialized mask. A midline cranial incision was then made and the skull was exposed and cleaned. A single hole (1 mm in diameter) was drilled in the skull in the lower left quadrant relative to bregma. After the horizontal skull position was confirmed, the cannula was placed according to bregma (-0.7 mm posterior, -1.2 mm lateral [left], and -2.0 mm ventral). G-ænial Bond and G-ænial Universal Flo (GC America, Inc., Alsip, IL, USA, G-Bond Unit Dose Kit Cat# 002,302 and G-ænial Universal Flo B1 Refill Cat# 004,207) were used to fix the cannula in place which was cured with LED light. Vetbond (3 M, St. Paul, MN, USA) was used to cloes the surgical incision. Gm8773 was injected (2 µL) using a Hamilton syringe connected to an infusion cannula with a 0.5 mm projection. Cannula placement was confirmed using NPY at a dose of 0.3 nmol/mouse (via 2 µL injections of 2.5 µg/µL NPY solution) injected in the early light phase. A cumulative food intake of ≥0.5 g in 3 h following ICV NPY injection indicated successful cannula placement.

#### METHOD DETAILS

**Isolation and differentiation of primary mouse white, beige, and brown adipocytes for both ribosome profiling and mass spectrometry**

Twenty female C57Bl6/J mice aged 7 weeks old were sacrificed by $CO_2$ asphyxiation followed by cervical dislocation. Interscapular and subscapular BAT was excised with dissecting scissors, being careful to avoid surrounding muscle and white adipose tissue. For white adipose tissue, subcutaneous (scWAT) depots were excised bilaterally. Fat pads were placed collectively in a 50 mL falcon containing ice-cold DMEM. DMEM was carefully decanted through a mesh filter, and the pads washed briefly in HBSS. Most of the HBSS was then decanted, before transferring each tube of fat pads to a single well of a 6-well culture plate. Dissecting scissors were then used to thoroughly mince each collection of pooled fat explants.

Digests were performed in the following solutions (pre-warmed to 37 ºC): 1) for scWAT, 40 mg Collagenase Type I with 80 mg Collagenase Type II in 28 mL HBSS and 12 mL 7.5% w/v BSA (Sigma); 2) for BAT, 80 mg Collagenase Type I with 80 mg Collagenase Type II in 28 mL HBSS and 12 mL 7.5% w/v BSA (Sigma). Digests were incubated at 37 ºC for 15-20 mins with shaking (225 rpm), resulting in complete dissociation of adipose tissue. Digests were then filtered through a 100 µm mesh (yellow, Corning) into 50 ml falcon tubes and allowed to sit on ice for 20 minutes. Visible fat floating as a layer was carefully aspirated. Working from the top, the supernatant was pipetted into a clean 50 ml Falcon tube, leaving behind the lowest 10 ml of digested material. This fraction is enriched for pre-adipocytes and was mixed 1:1 with 10% newborn calf serum (Gibco). Cells were spun down at 700 g for 15 minutes and resuspended in growth media consisting of: 1) for scWAT ("scWAT media"), DMEM plus 10% NCS with Pen/Strep, 4 mM Glutamax (Gibco), 150 µM sodium ascorbate, 30 nM insulin, and 1 µM rosiglitazone (Millipore Sigma, R2408); and 2) for BAT ("BAT media") DMEM plus 10% NCS with Pen/Strep, 4 mM Glutamax (Gibco), 150 µM sodium ascorbate, 4 nM insulin, and 2 nM T3 ( Millipore Sigma, T6397). Cells were spun again for 10 minutes and resuspended ready for plating in the following volumes (based on 2 ml of media for each well seeded): 1) for scWAT, a ratio of 5 mice in 38 mL of plating media with suspended cells, and 2) for BAT, a ratio of 5 mice in 25 mL of plating media with suspended cells. Cells were plated into 6-well dishes with 2 mL of cell suspension per well. Cells cultured as the "Beige" phenotype represent the scWAT cultures that were switched to the BAT media 24 hours after plating in the scWAT media. Media was changed daily for days 1-4 and every other day from days 4-8. On day 6, media was changed to phenol red free DMEM and rosiglitazone was removed from the scWAT cultures to allow wash out before harvest. All cultures were treated or harvested or both on day 8.

**Ribo-Seq and total RNA-Seq sample preparation from primary mouse white, beige, and brown adipocytes**

For sample collection of BAT/WAT/Beige cultures for Ribo-Seq and total mRNA-Seq used for smORF discovery, each culture was washed directly out of the incubators 3x with ice-cold PBS supplemented with 100µg/mL cycloheximide (CHX; Fisher Scientific, AAJ66004X). After the last wash, liquid nitrogen was gently ladled onto the surfaces of the cells and plates were stored at -80 °C prior to ribosome footprinting and preparation of sequencing libraries. Each biological replicate for Ribo-Seq in our experiments represents 11 wells of a 6-well dish leaving one well to be lysed with Trizol reagent (Thermo #15596026) for the bulk mRNA-Seq used int the *de novo* transcriptome assembly of the cells under analysis. Cells were lysed with 400 µl of ice-cold lysis buffer (20 mM Tris–HCl, pH 7.4, 150 mM NaCl, 5 mM MgCl2, 1% Triton X-100, with 1 mM DTT, 25 U $ml^{-1}$ Turbo DNase (Thermo Fisher, catalog no. AM2238) and 100 µg ml−1 CHX added fresh) was dripped onto the plate. Cells were incubated on ice in lysis buffer for 10 min with periodic vortexing and pipetting to disperse the cells. The lysate was then clarified by centrifugation at 15,000g for 10 min. Cell lysates were flash-frozen in liquid nitrogen and stored at −80 °C for up to 7 d before ribosome footprinting. For each cell type, ribosome footprinting was carried out by digesting 40-60 µg of RNA in 200-300 µL lysate with 0.375 U/$\mu g^{-1}$ RNase I (Lucigen, N6901K) for 50 min at room temperature. Digestion reactions were quenched with 200 U Superase-In RNase I inhibitor (Thermo Fisher, AM2694) on ice. Following digestion, monosomes were purified from small RNA fragments using MicroSpin S-400 HR columns (GE

Life Sciences), and ribosome protected RNA fragments (RPFs) were extracted by acid phenol chloroform and isopropanol precipitation. Sequencing libraries were prepared as in (McGlincy et al. 2017) with some modifications. First, the Ribo-Zero Mammalian Kit (Illumina) was used to deplete rRNA after RPF extraction and just prior to RPF size selection by gel extraction. Second, the Zymo clean & concentrator step after adaptor ligation is omitted and the reaction was carried over straight into reverse transcription. For the reverse transcription step to form cDNA, Episcript RT(Lucigen, ERT12910K) was used. Following reverse transcription, excess primer was degraded using Exonuclease I (Lucigen, X40520K) and the RNA templates were degraded using Hybridase (Lucigen, H39500). For the cDNA circularization step, CircLigase I (Lucigen, CL4111K) was used. PCR amplification was then carried out using Phusion Hot Start II High-Fidelity Master Mix (Thermo Fisher, F565L) for 9-12 cycles. The adapters and primers for library construction used were as follows: 3' adapter – 5'-/5phos/AGATCGGAAGAGCACACGTCTGAA/3ddC/-3'; RT primer – 5'-/5Phos/AGATCGGAAGAGCGTCGTGTAGGGAAAGAG/iSp18/GTGACTGGAGTTCAGACGTG TGCTC-3'; PCR forward primer – 5'-AATGATACGGCGACCACCGAGATCTACACTCTTTCCCTACACGACGCTC-3'; Illumina TruSeq Ribo Profile (Mammalian) Library Prep Kit index primers 1 – 12.

**Ribo-Seq and Bioinformatics Analysis**

We followed the methodology described in Martinez et al (Martinez et al., 2020) to generate custom open reading frame databases for microprotein discovery, with some key modifications. After trimming adapters, removal of mm10 rRNA and tRNA sequences, and alignment to the mm10 genome with STAR v2.53b (Dobin et al., 2013), instead of Cufflinks (Trapnell et al., 2010) Stringtie v2.1.4 (Trapnell et al., 2010) and MAPS v1.0 (Ma et al., 2018) were run using default parameters on combined alignments from total and mRNA RNA-Seq libraries from BAT, beige, and WAT tissues, followed by 3-frame translated using a custom script GTFtoFasta (see (Martinez et al., 2020) for details). The resulting 3-frame translated ORF databases were then scored for translation using RibORF (Ji, 2018) with the pipeline described by (Martinez et al., 2020), which briefly included shifting reads of each length to obtain base-pair resolution, and then scoring each candidate smORF using RibORF, keeping only highly scoring (score ≥ 0.7), short (≤ 150 aa), and novel (not overlapped with known coding regions in RefSeq, and a maximum blastp alignment evalue  to SwissProt (Bairoch and Apweiler, 1997). The resulting microproteins were then collated into a non-redundant table and were further annotated using HOMER ((Heinz et al., 2010); genomic location of their ORFs), and using PhyloCSF (Lin et al., 2011); conservation from a multi-way alignment of mammals) as described previously (Martinez et al., 2020). Candidate microproteins were then split into categories based on the genomic location of their ORF with respect to genomic features: uORFs exist upstream of known genes, non-uORFs are all other ORFs, iORFs specifically exist in intergenic regions, while dORFs exist downstream of known genes. These categories of ORFs were then further tested for expression in RNA-Seq datasets and visualized using read pileup tracks.

**Whole cell lysates ("proteomes") and secreted protein ("secretome") sample preparation from primary mouse white, beige, and brown adipocytes**

For whole cell proteome analysis, 1 mL of 8M urea (Sigma-Aldrich #U5128) in 100 mM ammonium bicarbonate (Sigma-Aldrich # A6141) (Urea lysis buffer, "ULB") was pipetted into each well of a 6-well dish of BAT-derived "brown" cells and scWAT-derived "white" cells that were differentiated with the protocol above where each well represents a single biological replicate. The proteome and secretome samples were prepared in parallel with the samples used for Ribo-Seq and total mRNA-Seq. Lysates were pipetted up and down to aid in lysis before collection. Additionally, parallel secretome samples were washed 3x with pre-warmed serum free phenol red free DMEM (Gibco # 21063029) (3x) followed by the addition of 500 uL of pre-warmed serum free phenol red free DMEM (Gibco #31053028). The cells were allowed to secrete proteins for 90 minutes before collection of the secretion media which was collected from each well into separate 2 mL LoBind eppendorf tubes and snap frozen in liquid nitrogen.

**Trypsin digestion and de-salting of differentiated brown and white adipocyte proteome and secretome samples**

For tryptic digestions, protein concentration estimates of each of the lysates were determined using BCA (Pierce #23225) and 100 µg of each sample was digested using a modified protocol suitable for S-trap (Protifi) digestion of lipid laden samples. The S-trap solubilization/lysis buffer is 5% sodium dodecyl sulfate (SDS, Sigma-Aldrich # 436143), 50 mM Triethylammonium bicarbonate buffer (TEAB, Sigma-Aldrich # T7408), 2 mM MgCl2 (Sigma-Aldrich # M8266), 1X HALT protease & phosphatase inhibitors (ThermoFisher # 78440) so each sample from the ULB extractions described above were brought to this final concentration prior to proceeding through the protocol. Once in the S-trap solubilization/lysis buffer, samples were reduced with 10 mM Tris(2-carboxyethyl)phosphine hydrochloride (TCEP, Sigma-Aldrich # C4706 ) for 30 minutes at room temperature with 600 RPM shaking on a thermomixer followed by alkylation with 40 mM iodoacetamide (Sigma-Aldrich # I1149 ) in the dark at room temperature with 600 RPM shaking on a thermomixer. 12% aqueous phosphoric acid was added at 1:10 ratio giving a concentration of ~1.2% phosphoric acid to neutralize iodoacetamide. Samples were vortexed and then spun down. S-trap binding buffer consisting of 90% methanol in 100 mM TEAB was added at 6x the volume of the sample, vortexed, then spun down. The acidified lysates with S-trap binding buffer were added to the 96-well S-traps columns and the plates were affixed to a plate vacuum manifold. Repeat additions of the sample were performed until all of each sample had passed through the columns. Columns were then washed with 200 µL S-trap binding buffer (3x). To remove lipids, columns were washed with 150 µL of a 50/50 mixture of chloroform and methanol (3x) followed by one more wash with S-trap binding buffer. The S-trap digestion plate was then moved to a clean LoBind collection plate and 5 µg of porcine trypsin (Promega, #V5113) suspended in 50 mM TEAB was added to each sample. The plates were loosely sealed with Teflon plate covers and incubated for 1 hr at 47 ºC. After digestion, 80 µL of 50 mM TEAB was added to all wells and the plates were centrifuged on top of the new collection plates at 1500 g for 2 min. An additional 80 µL 0.2% aqueous formic acid was added to all wells and centrifuged into the same collection plates at 1500 g for 2 min. Finally, 80 µL of 50% acetonitrile (ACN) containing 0.2% formic acid was added and centrifuged at 1500 g for 2 min for a final concentration of 10% v/v ACN. The elution plates were frozen at -80 ºC overnight with Teflon gaskets and wrapped in parafilm until holes were poked in the gaskets and the samples were lyophilized to dry. Samples were resuspended in 50 μL water with 0.1% formic acid (buffer A) supplemented with 50 fmol/μL Pierce retention time calibration (PRTC) peptides (Pierce #88321) and the plates were floated in a water bath sonicator for 5 minutes followed by 5 minutes on a thermomixer at 600 rpm prior to mass spectrometry analysis.

**Bulk mRNA-Seq analysis with next-generation sequencing in the DIO/lean tissue**

Mice were sacrificed and tissues of interest were extracted and kept at -80 °C until RNA extraction. Samples were thawed and homogenized in Trizol with one 5 mm stainless steel ball using a TissueLyser II (Qiagen). RNA was extracted using chloroform and a Qiagen RNeasy Mini Kit (#74004) following the manufacturer's instructions. RNA concentration and quality was measured on a Nanodrop and Bioanalyzer. Samples with a 260/280 ratio of > 1.8 and RIN > 8 were shipped on dry ice for sequencing at Covance Genomics Lab, Redmond, WA. cDNA libraries were prepared with TruSeq stranded mRNA kit (Illumina #20020594) and sequenced on an Illumina HiSeq 2000 using paired-end, 100 nucleotide reads with an average depth of 20 million reads per sample.

**Bioinformatics quantitation of NGS transcriptome data of bulk tissue (adipose depots and liver) mRNA DIO/lean**

Raw sequencing data was quality assessed using fastqc (https://www.bioinformatics.babraham.ac.uk/projects/fastqc/) (Wingett and Andrews, 2018). Reads were aligned to the mm10 reference genome using STAR v2.53b (Dobin et al., 2013) and reads were quantified using HOMER analyzeRepeats (Heinz et al., 2010) using the top expressed isoform as proxy for gene expression. Differential expression between DIO and lean mice was carried out using DESeq2 (Love et al., 2014) and genes with FDR < 0.05 and log2fold > 1 were identified as significantly changed. Gene pathway enrichment analysis was carried out using WebGestaltR (Liao et al., 2019) in ORA mode, using all genes as the background for testing. Candidate ORFs identified from Ribo-Seq were quantified across ORF exons analogously to annotated genes and differential ORF expression was tested with DESeq2 using a threshold of FDR < 0.05 and log2fold > 1. Principle component analysis (PCA) was carried out with the prcomp function and plotted in R. Ovals, were manually drawn to highlight specific groups of samples.

**Sample preparation and protein digestions including cleanup and desalting of tryptic peptides for DIO/lean young/old plasma**

For plasma protein digests for the DIO/lean young/old mouse experiments, 1 μL of plasma was diluted to 100 μL in water. 50 μL of this solution was combined with 50 μL of 0.2% PPS silent surfactant (Agilent # 400500) in 100 mM ammonium bicarbonate (Sigma-Aldrich # A6141) for a final concentration of 0.1% PPS and 50 mM ammonium bicarbonate. The remaining 50 μL was stored for BCA protein concentration estimates. The twice diluted plasma was incubated at 95 ºC for 5 min to facilitate protein denaturation and then reduced with the addition of 500 mM dithiothreitol (DTT) to a final concentration of 5 mM and incubation at 60 °C for 30 minutes. Reduced samples were then alkylated at 15 mM iodoacetamide at room temperature for 30 minutes in the dark. Alkylation was quenched by adding an additional aliquot of DTT to bring the final concentration to 10 mM. Each sample was digested with sequencing grade trypsin (Promega #V5113) at a dilution of 1:10 trypsin:protein with starting plasma concentrations estimated at 70 mg/mL such that 3.5 µg of trypsin was added to each sample. Digestion was allowed to proceed for 18 hours at 37 °C on a thermomixer set to 1300 RPM. The following day, digestion was quenched by the addition of 4.5 µL of 5 M HCL, which were allowed to incubate at room temperature for 1 hour to facilitate hydrolysis of the PPS surfactant. Samples were centrifuged at 15,000 g at 4 °C for 5 min to pellet insoluble material and precipitated PPS surfactant. Tryptic digests were desalted with MCX columns in 96-well format (Waters # 186001830BA) as follows. An MCX column plate was affixed to a vacuum manifold. Samples were added to the columns careful to avoid the precipitated PPS pellet at the bottom of the tube and then samples were allowed to sit for 10 min. Salts were washed with 1 mL 0.1% formic acid followed by a wash with 1 mL of 90% ACN / 10% water. The column plate was placed over the top of a new collection plate 600 µL 10% ammonium hydroxide in methanol was added to each column and allowed to sit for 30 min. Vacuum was used to pull the ammonium hydroxide / methanol mixture through to the new collection plates that were vacuum centrifuged to dryness. Samples were reconstituted in 20 µL buffer A supplemented with PRTC peptides were spiked in to a final concentration of 50 fmol/µL for DIA-MS analysis.

**Small protein sub-fractionation in mouse plasma**

For the quantification of plasma proteomes from the DIO/lean young/old experiment, an additional plasma protein physical fractionation step was employed with the goal of enriching for the small proteome. Normal lean plasma was enriched for small proteins in an effort to deeply identify microproteins in the circulation using the following four microprotein enrichment strategies. For this, 1 mL of lean mouse plasma (1 mL for each enrichment) was mixed with an equal volume of 8M guanidine-Cl (Pierce #24115), 0.2% Trifluoracetic acid (TFA, Pierce #28904), 1% β-mercaptoethanol (Fischer #PI35602) and filtered through a 0.45 µm filter. Samples were then enriched for peptides and small proteins, both known and novel, with 4 separate methods as follows. Two of the enrichments were performed with Agilent BondElut C18 SPE cartridges (Varian Product #12102028) using either triethylammonium formate buffer at pH 3.0 (TEAF) or 0.1% TFA, Pierce #28904 as the counter ion. TEAF was made by adding 23 mL of 88% formic acid to 1.9 L of $dH_2O$ and adjusting pH to 3.0 using triethylamine (Pierce #25108). The other two enrichments were performed with Agilent BondElut C8 SPE cartridges (Varian Product #12102029) also using either TEAF at pH 3.0 or 0.1% TFA. Eluted with 75% acetonitrile in counter ion. In all 4 cases, columns were prewet with 3 ml methanol followed by equilibration with 3 mL of either the TEAF at pH 3.0 solution or the 0.1% TFA solution. Samples were applied to the conditioned columns followed by a wash in either the TEAF at pH 3.0 solution or the 0.1% TFA solution. Samples were eluted in 75% acetonititrile and lyophilized prior to digestion. Each plasma fraction that had been enriched for the small proteome was digested with the plasma digest protocol in the section entitled “Sample preparation and protein digestions including cleanup and desalting of tryptic peptides for DIO/lean young/old plasma” with the peptides fractionated with disposable high pH reversed phase fractionation columns as described by the manufacturer (Thermo, #84868) as described in “Data-independent acquisition

(DIA) mass spectrometry for quantitative proteomics including spectral library generation with data-dependent acquisition (DDA)".

**Sample pooling and fractionation for mass spectrometry**

Sample pooling and queueing across all DDA and DIA experiments reported in this paper followed the strategy described in Fig. 2. Briefly, for each experiment, tryptic digested peptides from a representative batch of biological samples were pooled. These pools were used for both GPF-DIA acquisitions and high pH reversed phase fractionation for DDA to generate spectral libraries. For the experiment described in Fig. 3, separate pools were generated for proteome samples and for secretome samples. For the experiment described in Fig 5, subaliquots of all biological samples from all conditions were pooled. While an aliquot of each pool was reserved for GPF-DIA, these pools were additionally fractionated with disposable high pH reversed phase fractionation columns as described by the manufacturer (Thermo, #84868). Briefly, 80 µg of a pool of tryptic peptides was loaded onto the hydrophobic resin spin column and fractionally eluted using 8 buffers with increasing concentrations of acetonitrile as follows: 5%, 7.5%, 10%, 12.5%, 15%, 17.5%, 20.0%, and 50%. The peptide fractions were dried completely with vacuum centrifugation and resuspended in 15 µL of buffer A for further analysis with DDA.

**Data-dependent acquisition mass spectrometry (DDA-MS)**

Each fraction described in the previous section was injected into a Thermo Fusion Lumos attached to a Thermo EASY-nLC 1200. For DDA, tryptic digest fractions were separated on self-packed 30 cm columns packed with 1.8 µm ReproSil-Pur C18 silica beads (Dr. Maisch) inside of a 75 µm inner diameter fused silica capillary (#PF360 Self-Pack PicoFrit, New Objective). The 30 cm column was coiled inside of a Sonation PRSO-V1 column oven set to 55 °C prior to ionization into the MS. The HPLC was performed using 200 nl/min flow with solvent A as 0.1% formic acid in water and solvent B as 0.1% formic acid in 80% acetonitrile. For each injection, 3 µl (approximately 1 µg) was loaded and eluted with a linear gradient from 7% to 38% buffer B over 90 min. Following the linear separation, the system was ramped up to 75% buffer B over 5 min and finally set to 100% buffer B for 15 min, which was followed by re-equilibration to 2% buffer B prior to the subsequent injection. Data were acquired using DDA with a maximum 3 s cycle time configuration between MS1s and with 30 s dynamic exclusion. Precursor spectra were collected from 300–1650m/z at 60,000 resolution (AGC target of 5e5, max IIT of 20 ms). MS/MS were collected on +2H to +7H precursors achieving a minimum AGC of 2e3. MS/MS scans were collected at 30,000 resolution (AGC target of 1e5, max IIT of 47 ms) with an isolation width of 1.4m/z with a NCE of 27.

**Data-independent acquisition mass spectrometry (DIA-MS)**

For DIA-MS analysis, there are two methods involved in this study. There is the small window GPF-DIA-MS method (4m/z) and the larger-window DIA-MS method used for individual sample quantitation (8m/z). For GPF-DIA-MS, the Thermo Fusion Lumos was set to acquire six progressive GPF-DIA-MS acquisitions of the specific biological sample pool used in each comparison using 120,000 precursor resolution and 30,000 fragment resolution. Automatic gain control (AGC) target was set to 4e5 with the fragment maximum ion inject time (IIT) set to 60 ms. The NCE was set to 33 and +2H was assumed as the default charge state. For the GPF-

DIA-MS acquisitions, we chose to use 4m/z precursor isolation windows in a staggered-window pattern with the following optimized window placements ranges: 398.4 to 502.5m/z, 498.5 to 602.5m/z, 598.5 to 702.6m/z, 698.6 to 802.6m/z, 798.6 to 902.7m/z, and 898.7 to 1002.7m/z (Amodei et al., 2019). To quantify the proteomes of individual samples, we used single-injection DIA-MS acquisitions (120,000 precursor resolution, 15,000 fragment resolution, AGC target of 4e5, fragment max IIT of 20 ms) with 8m/z overlapping precursor isolation windows in a staggered-window pattern with optimized window placements over a range of 396.4 to 1004.7m/z.

#### Bioinformatics for proteomics

##### *DDA data processing, database search, and spectral library creation for high pH reversed phase fractionated pools*

All Thermo RAW files were converted to mzXML format using ProteoWizard (version 3.0.9974) (Chambers et al., 2012). DDA-based peptide identifications were derived by searching mzXML files against a custom mouse proteome database that included the canonical, reviewed *Mus musculus* database downloaded from uniprot.org amended with the 3,877 microprotein sequences compiled from the Ribo-Seq smORF discovery work performed on the primary mouse BAT/WAT/Beige cells (Table S1). Peptide identifications were derived by concatenated target-decoy searching using Comet (version 2018.01 rev. 0)(Eng et al., 2013) with a modified "high-high" parameters set. In brief, this included variable modifications of up to three methionines per peptide, peptide n-terminal pyro-glutamate formation on glutamine, and protein n-terminal acetylation as well as fixed cysteine carbamidomethylation. Additionally, the "clip_nterm_methionine" parameter was turned on. Fully tryptic searches were performed with a 20 ppm precursor tolerance and a 0.02 Da fragment tolerance permitting up to two missed cleavages. High-pH reversed-phase fractions were combined and search results were filtered to a 1% peptide spectrum match (PSM) FDR using PeptideProphet (Keller et al., 2002) from the Trans-Proteomic Pipeline (TPP version 5.1.0) (Deutsch et al., 2015). For the proteome and secretome experiments in Fig. 3, a combined spectral library from both compartments was used for the subsequent DIA-MS. For the plasma proteome work in Fig. 5, a combined spectral library was generated from searches against all of the high pH reveresed phase fractions of the experimental samples themsleves along with all of the small protein enriched lean plasma samples that were also digested and fractionated with high pH reversed phase fractionation. The spectral libraries generated for Fig. 3 and Fig 5 were not combined such that the Fig. 3 spectral library was used in the Fig. 3 DIA-MS and the Fig. 5 spectral library was used in the Fig. 5 DIA-MS. Search results were imported into Sklyine such that the peptide prophet probability that correlated to a 1% PSM-level FDR was added as the "cut-off score" in Skyline where the search results were converted to .blib format for use in DIA-MS workflows to follow. Fragment spectra from key DDA peptides (VFCHQANDVHIYQTQVVMTNTLETSSGK++++ and MINLLMQHQR++) were visually validated with deep-learning fragmentation pattern predictions generated by Prosit (Gessulat et al., 2019) using model version: "Prosit_2020_intensity_hcd".

***DIA data processing for DIO/lean adipose tissue atlas, DIO/lean young/old plasma proteome profiling, and BAT/WAT/Beige proteome and secretome analysis***

All Thermo RAW files were converted to mzML format and overlap demultiplexed with 10 ppm accuracy after peak picking using ProteoWizard (version 3.0.9974). The GPF-DIA-MS injections for each experimental comparison were searched against the experiment-specific spectral library (.blib from DDA search above) to derive peptide identifications with accurate chromatographic retention time information for each peptide using EncyclopeDIA (version 0.9.0), which was configured with the default settings of 10 ppm precursor, fragment, and library tolerances. Both B and Y ions were allowed to be considered with trypsin digestion assumed in the EncyclopeDIA settings set. Following the protocol outlined in (Searle et al., 2018, 2020), all EncyclopeDIA searches were performed with built in target/decoy (Elias and Gygi, 2007) peptide-level FDR estimation using Percolator 3.1 (Käll et al., 2007; The et al., 2016). Proteins were then parsimoniously grouped and filtered to a 1% protein-level FDR. The EncyclopeDIA .elib report files were imported into Skyline, along with the individual DIA-MS raw files for data visualization and group comparisons.

**QPCR for Gm8773 mRNA in Mouse Tissue Extracts**

***Tissue Harvest***

Female or male mice (n = 6) at 8–9 weeks of age from each strain were sacrificed by halothane inhalation (Halocarbon Laboratories; River Edge, NJ) at lights on (ZT0). All tissues were from male mice except for female-reproductive tissues. Mice were exsanguinated via the vena cava with whole tissues collected and snap frozen in liquid nitrogen. Tissues were isolated from appropriate anatomical locations according to established methods or (Iwaki et al., 2001). Brown adipose was collected from the dorsal interscapular depression and any surrounding white fat or connective tissue was removed. White adipose was collected from the epididymal area. Skeletal muscle from thequadriceps was isolated from both femurs, and bone marrow was removed from the remaining bone by flushing with PBS. The enteric tract, including duodenum, jejunum, ileum, colon, and rectum was sectioned and flushed with PBS, and the intestinal mucosa scraped and collected. The brain was sectioned into hypothalamus, pituitary, brain stem, cerebrum, olfactory bulb, cerebellum, and corpus striatum, which was the remaining brain after the other sections had been removed (Paxinos and Franklin, 2019). An 18G needle was used to push the upper spinal cord out of the spinal column . Both uterine horns were collected. Dorsal skin was shaved prior to collection. The pancreas was prepared immeditedly due to its high RNase content by collection directly into the RNA-isolation reagent. The rest of the snap-frozen tissues were stored at −80°C until RNA extraction.

***RNA Isolation and cDNA Preparation***

RNAStat60 (TelTest; Friendswood, TX) was used to extract RNA according to the manufacturer's directions with a few tissue-specific modifications to aid in the mechanical disruption of some organs prior to RNA isolation. A Bessmann pulverizer (Fisher Scientific) was used to crush skeletal muscle, bone, and skin were crushed into a powder. Whole livers were also crushed and homogenized into powder to minimize differences lobe-to-lobe mRNA expression . Total RNA was pooled in equal quantities for each tissue (n = 6). RNA pools from male mice were used for all tissue analyses except those of the ovary and uterus. DNase treatment with Ambion's Turbo DNA-free kit (Austin, TX) was used to eliminate genomic, except for pancreas and seminal vesicle RNA, which were sensitive to degradation. Preparation of cDNA for QPCR assays was performed as previously described (Bookout and Mangelsdorf, 2003) with the following changes: 2.4 µg of RNA was first treated with 2U DNase I and 4.2 mM MgCl2 in a final volume of 40 µl. The reverse-transcription reaction was carried out in 100 µl final volume. Following cDNA synthesis, DEPC-H2O was added to increase the sample volume to 300 µl.

***QPCR***

Extracted tissue mRNA levels from extracts described above were measured using an ABI 7900HT Sequence Detection System as described previously (Bookout et al., 2006a, 2006b). Prior to pooling RNA samples for Gm8773 expression analysis, individual tissue samples were assayed for certain difficult-to-dissect tissues to ensure their fidelity. Tissue-specific marker expression was confirmed using the SYBR Green ΔΔCt method as described (Bookout et al., 2006a) with the primers summarized in the Key Resources Table. Individual RNAs were compared to known, commercially available samples (Ambion, BD Clontech, Harlan Bioproducts, Stratagene).

Analysis of Gm8773 mRNA expression was performed using the TaqMan-based efficiency-corrected ΔCt assay with 10 ng cDNA per reaction for 50 cycles (Bookout et al., 2006a). Nuclear receptor mRNAs with cycle times ≥ 34 were determined to be below detection. Primer concentrations were 75 nM for 18S rRNA and 300 nM for Gm8773 primers (primers listed in Table Key Resources Table), The primer/probe sets for Gm8773 expression were validated as described (Fu et al., 2005). Universal cDNA standards generated from mouse RNA (BD Clontech) were used for analysis of all receptors except CAR, FXRβ, PXR, SHP, DAX-1, ERβ, LRH-1, PNR, SF1, and TLX, which were too limited in expression to use the universal RNA set. For these receptors tissue-specific total RNA standards were used from liver, ovary, eye, adrenal, and whole brain, as appropriate.

ABI instrument software SDS2.1 was used to analyze all QPCR data. Baseline values of amplification plots were automatically set and threshold values kept constant to obtain normalized cycle times and linear regression data. Individual PCR efficiencies were were calculated from the slope of the resulting standard curves as reported previously (Fu et al., 2005). Normalized mRNA levels are expressed as arbitrary units and were obtained by dividing the averaged, efficiency-corrected values for Gm8773 mRNA expression by that for 18S RNA expression for each sample. The resulting values were multiplied by 105 for graphical representation and plotted ± standard deviation (S.D.) from triplicate sample wells.

***In situ* hybridization and visualization of Gm8773 transcript in the mouse brain**

Formalin-fixed, paraffin-embedded (FFPE) blocks containing mouse brain were sectioned at 5 µm onto Fisher SuperFrost Plus glass (Fisher Scientific, Hampton, NH, USA). Sections were hybridized with a rodent-specific probe to detect mouse Gm8773 mRNA (#542179, Advanced

Cell Diagnostics, Newark, CA) on a Ventana Discovery ULTRA (Ventana Medical Systems, Tucson, AZ, USA), and amplified/stained using an RNAscope RED kit (Advanced Cell Diagnostics), as described previously (Hultman et al., 2019). In this assay we used our standard pre-treatment conditions (24 min target retrieval, 12 min protease) to probe Gm8773 while maintaining CNS morphometry. After ISH, all slides were counterstained with hematoxylin and bluing, and were coverslipped using EcoMount (BioCare, Pacheco, CA) All sections were imaged at 20x on an AxioScan.Z1 (Zeiss, Jena, Germany). Post-hoc processing matched brightness/contrast across all slides, and images were compiled in Adobe Illustrator (Adobe Inc., San Jose, CA) for presentation.

#### Recombinant expression and purification of Gm8773 protein product

##### *Plasmids*

For recombinant expression, synthetic sequences encoding mouse Gm8773 (Uniprot Q3UQ24 D25-I132) was cloned into a mammalian expression vector encoding a N-terminal HSA fusion tags with expression under the control of the CMV promoter. The native signal peptide as predicted by SignalP was replaced with the CD33 signal peptide, yielding a final sequence of D25-I132. Sigma GenElute Endofree maxi prep kits (prod# NA0410-1KT) were used to generate higher amounts of plasmid DNA for scale up production efforts.

##### *Generation of recombinant proteins in transient mammalian host systems*

For transient protein expression, the Gibco Expi293 Expression system (prod# A14525) was used, following the vendor's recommended protocol. Cells were grown at 37°C in Expi293 medium™ (prod# A14351-01) at 5-8%CO2, 85% humidity, and shaking, with a maximum passage number of 30. The day prior to transfection, cells were split to 2e6 cells/ml. The day of transfection, cells were counted and transferred to a 1.6 L flask (Thomson, 931113) at 2.9e6 cells/ml. Plasmid DNA was combined with Expifectamine™ 293 Transfection Reagent following the vendor's suggested concentration and ratio. Cultivation was continued for 18 - 20 h at 37°C, 5-8%CO2, 85% humidity, and shaking, at which point the cultures were supplemented with ExpiFectamine™ 293 Transfection Enhancer 1 and Enhancer 2. After a total of 5 days post-transfection, the cell supernatants were collected by centrifugation and filtration (0.2 uM Millipore S2GPU11RE) and stored at 4°C.

##### *Purification of Gm8773*

900mL of filtered supernatant was captured on 2x5ml Ni excel columns at 5ml/min, washed with 5 column volumes (CV) of PBS pH7.2 and eluted in 2.5CV of PBS pH 7.2, 300mM Imidazole, 300mM L-Arginine. Selected fractions were pooled for a total of 256mg and digested with 2.5mg HRV 3C for a 1:100 dilution overnight at 4C. After cleavage the pooled fractions were applied to a Superdex 75 (Millipore Sigma GE17-5174-01) in PBS pH 7.2 + 200mM L-Arginine Buffer to separate the HSA tag from the now untagged protein. Selected pooled fractions were characterized by SDS-PAGE, analytical size exclusion and LC-MS.

***Analysis of Gm8773***

For analysis of deglycoslated protein 20ug of protein was incubated with 0.5uL of SialExo enzyme (Genovis, # G1-SM1-020) for 1 hour at 37C. SDS-PAGE analysis was performed using Bolt 4-12% Bis-Tris gels (Invitrogen NW04122BOX) and stained using the eStain system (Genscript L00753). LC/MS analysis was performed using an Agilent 1290 Infinity II UHPLC system coupled to an Agilent 6545 Q-TOF. LC separations were performed on a Zorbax 300-Diphenyl column (Agilent, 1.8 µm, 300 Å) with a flow rate of 0.4 ml/min used with a solvent gradient of 30% to 70% B in 6 minutes. Solvent A was 0.1% (v/v) formic acid in water and the composition of solvent B was 0.1% (v/v) formic acid in 100% acetonitrile. The mass spectrometer was operated in positive ion mode with a full-scan MS spectra from 400 to 3,200 m/z. Analytical size exclusion was performed using an Agilent UPLC HP1290 system using gel filtration standards (BioRad 1511901). Separations were performed on a Waters BEH column, 1.7 µm, 200 Å, 4.6 mm ID x 150 mm L (Waters, # 186005225) with a flow rate of 0.35 ml/minute in 10 minutes. The mobile phase was 20 mM Phosphate, 150 mM NaCl, 2% Isopropanol, pH 7.

**Acute food intake effects of ICV Gm8773 injections in DIO mice**

Acute effects on food intake were monitored in DIO mice with the ICV-cannulations described above. DIO mice were which were male C57BL/6J mice purchased from Jackson Laboratories with obesity induced in-house via continuous feeding with high fat diet (45% HFD Research Diets: D12451i) from age 10 weeks on. DIO mice were cannulated for ICV injections at age 52 weeks. Food intake was monitored using a fully automated human-intereference-free food-intake monitoring system that allows for continuous FI monitoring over time (BioDAQ, Research Diets, Inc., New Brunswick, NJ, USA). Animals were randomized into groups according to body weight and 48 hour pre FI prior to the ICV injections (n = 8). Animals each received one ICV injection of vehicle or test compound (0.495 nmol/mouse, dose volume 3µl). Compounds were dosed to four-hour-fasted mice just prior to lights out, and food intake data were collected for 24 h post dosing.

#### QUANTIFICATION AND STATISTICAL ANALYSIS

All data were analyzed with GraphPad Prism 9.0.1. Bar plots in Figure 3 represent the DIA-MS-based summed fragment ion intensities of each peptide for each protein/microprotein entry with at least one peptide per ORF/smORF where statistical comparisons were calculated using a two-tailed Student's t-test where values represent the means ± SD and asterisks indicate * = p-value < 0.05, ** = p-value < 0.01, and *** = p-value < 0.001. Values in Fig. 5F represent the same DIA-MS-based summed fragment ion intensity depicting the mean ± SD with statistics performed using a one-way ANOVA where **** = p-value < 0.0001. Lastly, the ICV-based food intake values in Fig. 6H with statistics performed with a two-way ANOVA with significant time by treatment interaction where **** = p-value < 0.0001.

#### KEY RESOURCES TABLE

| REAGENT or RESOURCE | SOURCE | IDENTIFIER |
|---|---|---|
| Antibodies | | |
| none | | |
| Bacterial and virus strains | | |
| none | | |
| Biological samples | | |
| Healthy C57BL/6J female mouse brown and subcutaneous white adipose tissue (7 week old) | The Jackson Laboratory | JAX: 000664 |
| Diet-induced obese 27-week old C57BL/6J male mouse liver, brown adipose tissue (BAT), epididymal white adipose tissue ("eWAT"), subcutaneous white adipose tissue (scWAT), retroperitoneal fat ("Retro Fat") and mesenteric fat ("Mesen Fat") fed high-fat diet for 21 weeks along with age matched counterparts fed chow | The Jackson Laboratory | JAX: 000664 |
| Aged (41-weeks) diet-induced obese (fed *ad lib* 45% HFD from age 10 weeks) C57BL/6J male mouse plasma | The Jackson Laboratory | JAX: 000664 |
| Aged (41-weeks) lean C57BL/6J male mouse plasma | The Jackson Laboratory | JAX: 000664 |
| Young (26-weeks) diet-induced obese (fed *ad lib* 45% HFD from age 10 weeks) C57BL/6J male mouse plasma | The Jackson Laboratory | JAX: 000664 |
| Young (26-weeks) lean mouse plasma | The Jackson Laboratory | JAX: 000664 |
| Male and female C57BL/6J mouse tissues RNA extract bank for qPCR | The Jackson Laboratory | JAX: 000664 |
| Chemicals, peptides, and recombinant proteins | | |
| DMEM | ThermoFisher / Gibco | Cat# 10566016 |
| HBSS | ThermoFisher / Gibco | Cat# 14025134 |
| Collagenase Type I | ThermoFisher / Gibco | Cat# 17100017 |
| Collagenase Type II | ThermoFisher / Gibco | Cat# 17101015 |
| BSA | Millipore Sigma | Cat# A9418 |
| Newborn calf serum | ThermoFisher / Gibco | Cat# 16010167 |
| Glutamax | ThermoFisher / Gibco | Cat# 35050079 |
| Sodium ascorbate | Millipore Sigma | Cat# A7631 |
| Insulin | ThermoFisher / Gibco | Cat# 12585014 |
| Rosiglitazone | Millipore Sigma | Cat# R2408 |
| T3 | Millipore Sigma | Cat# T6397 |
| Cycloheximide | Fisher Scientific | Cat# AAJ66004X |
| Trizol | ThermoFisher | Cat# 15596026 |

| | | |
|---|---|---|
| Triton X-100 | ThermoFisher | Cat# A16046.AP |
| Turbo DNase | ThermoFisher | Cat# AM2238 |
| RNase I | Lucigen | Cat# N6901K |
| Superase-In RNase I inhibitor | ThermoFisher | Cat# AM2694 |
| Episcript RT | Lucigen | Cat# ERT12910K |
| Exonuclease I | Lucigen | Cat# X40520K |
| Hybridase | Lucigen | Cat# H39500 |
| CircLigase I | Lucigen | Cat# CL4111K |
| Phusion Hot Start II High-Fidelity Master Mix | ThermoFisher | Cat# F565L |
| Urea | Sigma-Aldrich | Cat# U5128 |
| ammonium bicarbonate | Sigma-Aldrich | Cat# A6141 |
| DMEM, high glucose, no glutamine, no phenol red | ThermoFisher / Gibco | Cat# 31053028 |
| sodium dodecyl sulfate | Sigma-Aldrich | Cat# 436143 |
| Triethylammonium bicarbonate buffer | Sigma-Aldrich | Cat# T7408 |
| MgCl2 | Sigma-Aldrich | Cat# M8266 |
| HALT protease & phosphatase inhibitors | ThermoFisher | Cat# 78440 |
| Tris(2-carboxyethyl)phosphine hydrochloride | Sigma-Aldrich | Cat# C4706 |
| Iodoacetamide | Sigma-Aldrich | Cat# I1149 |
| porcine trypsin | Promega | Cat# V5113 |
| Pierce retention time calibration (PRTC) peptides | Pierce | Cat# 88321 |
| Rodent diet with 60 kcal% from fat | Research Diets | Cat# D12492i |
| Rodent diet with 10 kcal% from fat | Research Diets | Cat# D12450Bi |
| PicoLab Rodent Diet 20 | LabDiet | Cat# 5053 |
| Rodent diet with 45 kcal% from fat | Research Diets | Cat# D12451i |
| PPS silent surfactant | Agilent | Cat# 400500 |
| 8M guanidine-Cl | Pierce | Cat# 24115 |
| Trifluoracetic acid | Pierce | Cat# 28904 |
| β-mercaptoethanol | Fischer Scientific | Cat# PI35602 |
| Triethylamine | Pierce | Cat# 25108 |
| Standard rodent diet | Harlan Teklad | Cat# 7001 |
| RNAStat60 | Tel-Test, Inc. | Cat# RNA STAT-60 |
| RNAscope rodent-specific probe for mouse Gm8773 mRNA | Advanced Cell Diagnostics | Cat# 542179 |
| RNAscope RED kit | Advanced Cell Diagnostics | Cat# 322350 |
| Expi293 medium | ThermoFisher / Gibco | Cat# A14351-01 |
| Expifectamine™ 293 Transfection Reagent | ThermoFisher / Gibco | Cat# A14524 |
| SialExo enzyme | Genovis | Cat# G1-SM1-020 |
| gel filtration standards | BioRad | Cat# 1511901 |
| Critical commercial assays | | |
| Ribo-Zero Mammalian Kit | Illumina | Cat# 20040526 |
| TruSeq Ribo Profile (Mammalian) Library Prep Kit index primers 1 – 12. | Illumina | Cat# RPYSC12116 |
| BCA Protein Assay Kit | ThermoFisher / Pierce | Cat# 23225 |
| RNeasy Mini Kit | Qiagen | Cat# 74004 |
| TruSeq stranded mRNA kit | Illumina | Cat# 20020594 |
| Deposited data | | |

| | | |
|---|---|---|
| GEO Superseries ID of all raw and analyzed data | This paper | GEO: GSE198109 |
| Raw and analyzed Ribo-Seq data | This paper | GEO: GSE197909 |
| Raw and analyzed mRNA-Seq data of primary metabolic cells | This paper | GEO: GSE198107 |
| Tissue mRNA-Seq data of DIO and lean mice | doi: https://doi.org/10.1101/2021.06.25.449953 | GEO: GSE185466 |
| Mouse Primary brown, beige, and white whole cell lysate proteome data (released to ProteomeXchange upon publication) | This paper | MassIVE: MSV000089022 |
| Mouse Primary brown, beige, and white secretome proteome data (released to ProteomeXchange upon publication) | This paper | MassIVE MSV000089023 |
| Mouse plasma proteome data from DIO/lean young/old mice (released to ProteomeXchange upon publication) | This paper | MassIVE MSV000089021 |
| Experimental models: Cell lines | | |
| none | | |
| Experimental models: Organisms/strains | | |
| Mouse: C57Bl6/J 7 week old female | The Jackson Laboratory | JAX: 000664 |
| Mouse: C57Bl6/J 7 week old male | The Jackson Laboratory | JAX: 000664 |
| Oligonucleotides | | |
| Ribo-Seq Library Construction - 3' adapter – 5'-/5phos/AGATCGGAAGAGCACACGTCTGAA/3ddC/-3' | Martinez et al., 2020 | N/A |
| Ribo-Seq Library Construction - RT primer – 5'-/5Phos/AGATCGGAAGAGCGTCGTGTAGGGAAAGAG/iSp18/GTGACTGGAGTTCAGACGTGTGCTC-3' | Martinez et al., 2020 | N/A |
| Ribo-Seq Library Construction - PCR forward primer – 5'-AATGATACGGCGACCACCGAGATCTACACTCTTTCCCTACACGACGCTC-3' | Martinez et al., 2020 | N/A |
| 18s General SYBR Green qPCR primer – 5'–ACCGCAGCTAGGAATAATGGA-3' | Bookout et al., 2006a | N/A |
| 18s General SYBR Green qPCR primer – 5'–GCCTCAGTTCCGAAAACCA-3' | Bookout et al., 2006a | N/A |
| Gm8773 (NR_033499) SYBR Green qPCR primer – 5'-GCGTGGCCACCCACTCT | This paper | N/A |
| Gm8773 (NR_033499) SYBR Green qPCR primer – 5' – GCAGGACCTCGCTCCTTTTC-3' | This paper | N/A |
| Software and algorithms | | |

| | | |
|---|---|---|
| STAR v2.53b | Dobin et al., 2013 | https://github.com/alexdobin/STAR |
| Cufflinks | Trapnell et al., 2010 | https://github.com/cole-trapnell-lab/cufflinks |
| Stringtie v2.1.4 | Trapnell et al., 2010 | https://github.com/skovaka/stringtie2 |
| MAPS v1.0 | Ma et al., 2018 | https://bitbucket.org/shokhirev/maps/src/master/ |
| GTFtoFasta | Martinez et al., 2020 | N/A |
| RibORF | Ji, 2018 | https://github.com/zhejilab/RibORF |
| HOMER | Heinz et al., 2010 | http://homer.ucsd.edu/homer/ |
| PhyloCSF | Lin et al., 2011 | https://github.com/mlin/PhyloCSF |
| fastqc | Wingett and Andrews, 2018 | https://www.bioinformatics.babraham.ac.uk/projects/fastqc/ |
| DESeq2 | Love et al., 2014 | https://bioconductor.org/packages/release/bioc/html/DESeq2.html |
| WebGestaltR | Liao et al., 2019 | https://github.com/bzhanglab/WebGestaltR |
| ProteoWizard version 3.0.9974 | Chambers et al., 2012 | https://proteowizard.sourceforge.io/ |
| Comet (version 2018.01 rev. 0) | Eng et al., 2013 | http://comet-ms.sourceforge.net/ |
| Trans-Proteomic Pipeline (TPP version 5.1.0) | Deutsch et al., 2015 | https://sourceforge.net/projects/sashimi/files/Trans-Proteomic%20Pipeline%20%28TPP%29/ |
| Skyline (version 21.1.0.146) | MacLean et al., 2010 | https://skyline.ms/project/home/software/Skyline/begin.view |
| Prosit | Gessulat et al., 2019 | https://github.com/kusterlab/prosit/ |
| EncyclopeDIA (version 0.9.0) | Searle et al., 2018, 2020 | https://bitbucket.org/searleb/encyclopedia/wiki/Home |
| Percolator 3.1 | Käll et al., 2007; The et al., 2016 | https://github.com/percolator |

| Other | | |
|---|---|---|
| 100 µm cell strainer | Corning | Cat# 431752 |
| MicroSpin S-400 HR columns | Millipore Sigma | Cat# GE27-5140-01 |
| S-trap | Protifi | N/A |
| Li Heparin Microtainer tubes | Fisher Scientific | Cat# 365965 |
| MCX columns in 96-well format | Waters | Cat# 186001830BA |
| BondElut C18 SPE cartridges | Agilent / Varian | Cat# 12102028 |
| High pH Reversed-Phase Peptide Fractionation Kit | ThermoFisher | Cat# 84868 |
| 1.9 µm ReproSil-Pur C18 silica beads | Dr. Maisch | Cat# r119.b9 |
| 75 µm inner diameter fused silica capillary, Self-Pack PicoFrit | New Objective | Cat# PF360 |
| Ambion Turbo DNA-free kit (Austin, TX | Fisher Scientific | Cat# AM1907 |
| Sigma GenElute Endofree maxi prep kits (prod# NA0410-1KT | Millipore Sigma | Cat# NA0410-1KT |
| ExpiFectamine™ 293 Transfection Kit | ThermoFisher | Cat# A14525 |
| Superdex 75 | Millipore Sigma | Cat# GE17-5174-01 |
| eStain™ L1 Protein Staining System | Genscript | Cat# L00753 |
| Bolt 4-12% Bis-Tris gels | ThermoFisher | Cat# NW04122BOX |
| Zorbax 300-Diphenyl column (1.8 µm, C8) | Agilent | Cat# 863750-944 |
| Waters BEH column, 1.7 µm, 200 Å, 4.6 mm ID x 150 mm L | Waters | Cat# 186005225 |
| 26-gauge-guided cannula | PlasticsOne | Cat# C315GS-4/SPC |
| G-ænial Bond and G-ænial Universal Flo:<br>G-Bond Unit Dose Kit<br>G-ænial Universal Flo B1 Refill | GC America | Cat# 002302<br>Cat# 004207 |
| Vetbond | 3M | Cat# 70200742529 |

### Fig. S1

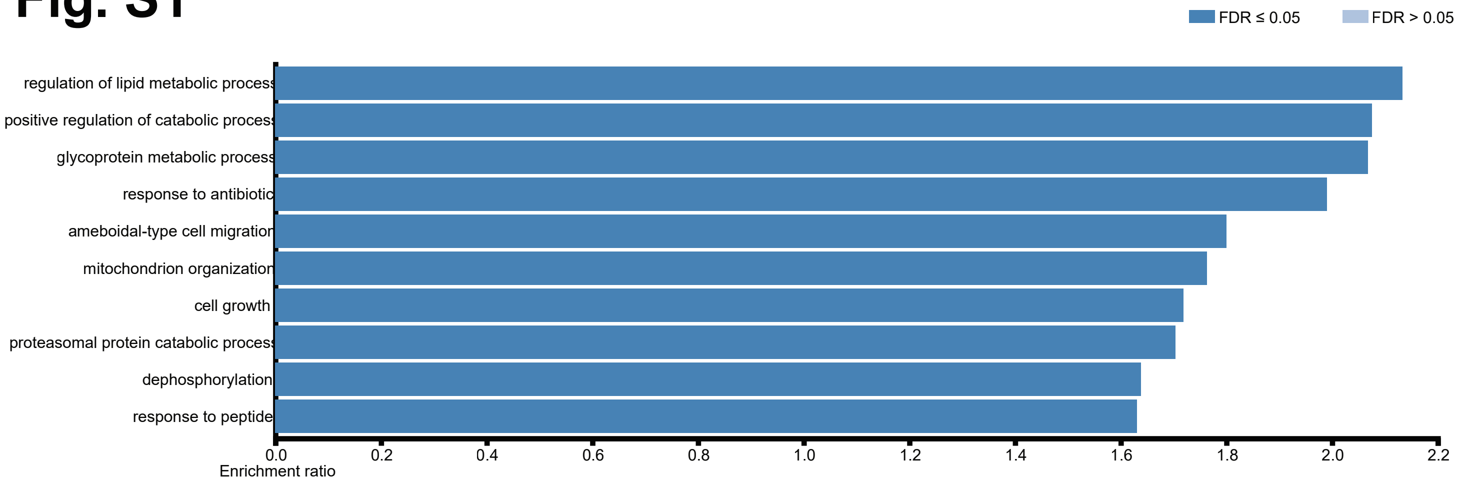

FDR ≤ 0.05
FDR > 0.05
regulation of lipid metabolic process
positive regulation of catabolic process
glycoprotein metabolic process
response to antibiotic
ameboidal-type cell migration
mitochondrion organization
cell growth
proteasomal protein catabolic process
dephosphorylation
response to peptide
0.0
0.2
0.4
0.6
0.8
1.0
1.2
1.4
1.6
1.8
2.0
2.2
Enrichment ratio

**Fig. S1. Gene ontology (GO) biological process analysis of uORF-containing genes.** GO enrichment analysis on the annotated genes containing uORFs listed in Table S1 (Ashburner et al., 2000). The top 10 redundancy-filtered pathways are shown.

Fig. S2

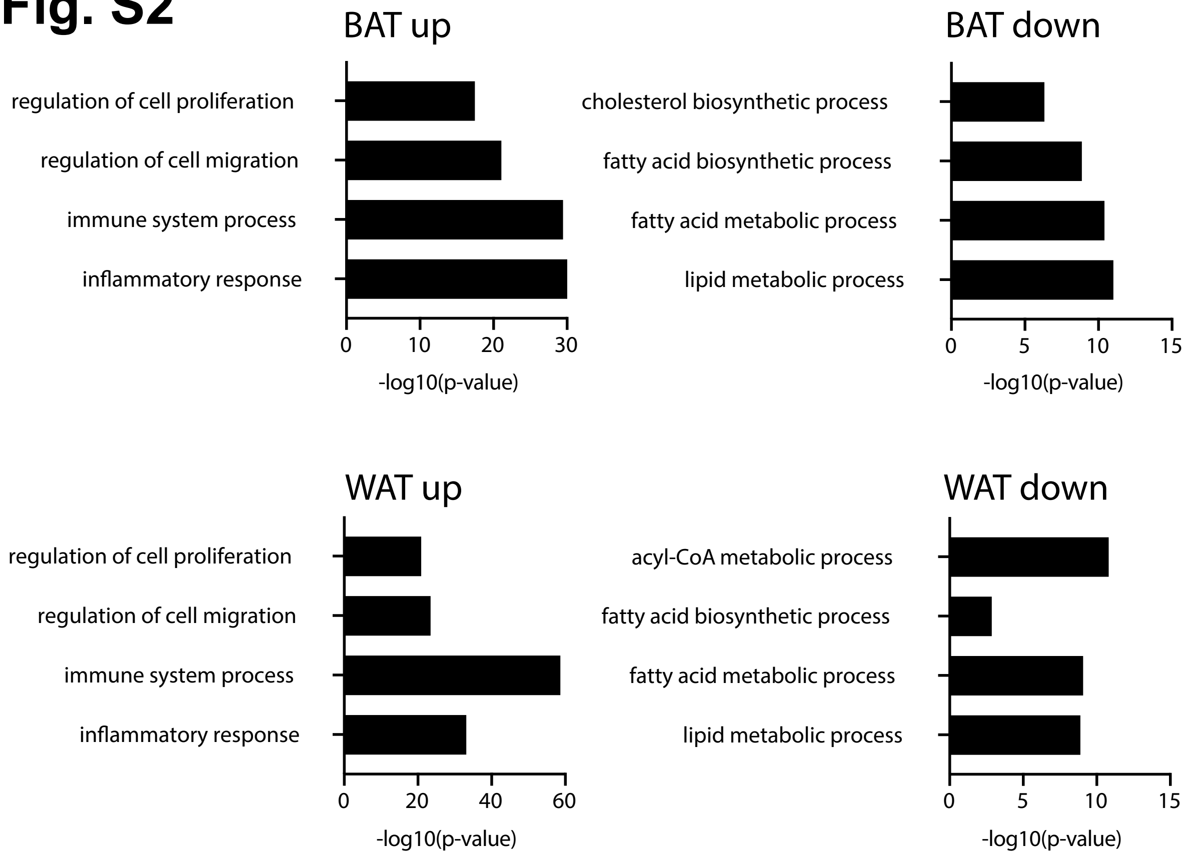
BAT up
regulation of cell proliferation
regulation of cell migration
immune system process
inflammatory response
0
10
20
30
-log10(p-value)
BAT down
cholesterol biosynthetic process
fatty acid biosynthetic process
fatty acid metabolic process
lipid metabolic process
0
5
10
15
-log10(p-value)
WAT up
regulation of cell proliferation
regulation of cell migration
immune system process
inflammatory response
0
20
40
60
-log10(p-value)
WAT down
acyl-CoA metabolic process
fatty acid biosynthetic process
fatty acid metabolic process
lipid metabolic process
0
5
10
15
-log10(p-value)

**Fig. S2. Gene ontology (GO) biological process analysis of bulk RNA-Seq-based transcriptomics expression of smORF genes up- and down- regulated in both BAT and epididymal WAT ("eWAT") tissue depots from both DIO and lean mice.** smORF coordinates listed in Table S1 were used to extract DIO vs. lean expression levels of uORF-containing smORFs. Depicted in Fig. S2 are up- and down-regulated GO-enriched processes where changes in RNA expression for adipose protein-coding smORFs induced by DIO in various tissues (padj $< 0.05$ and $|\log 2 \text{ fold change}| \geq 1$).

### Fig. S3

## A

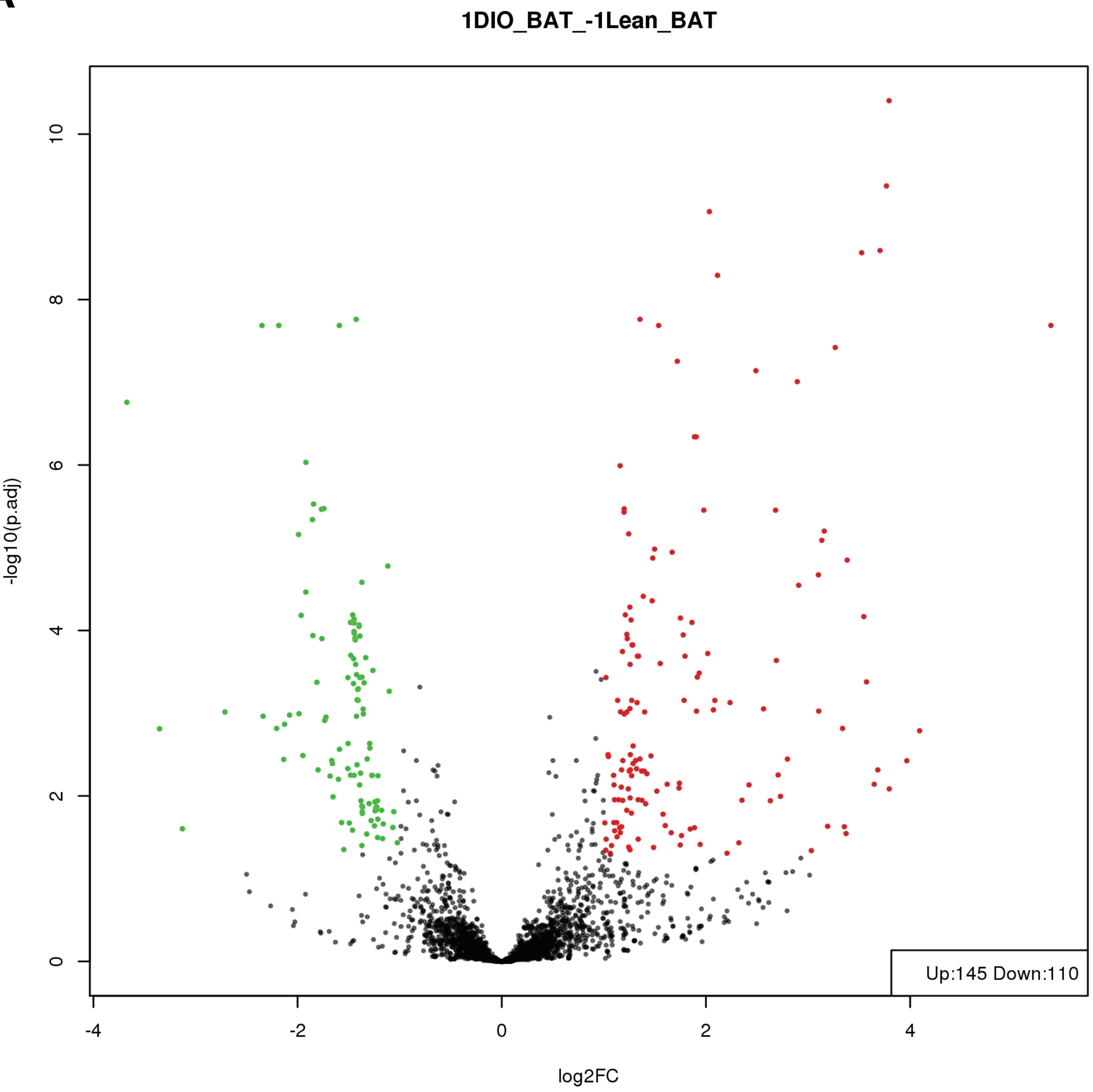
1DIO_BAT_-1Lean_BAT
-log10(p.adj)
log2FC
Up:145 Down:110
0
2
4
6
8
10
-4
-2
0
2
4

Fig. S3

**B**

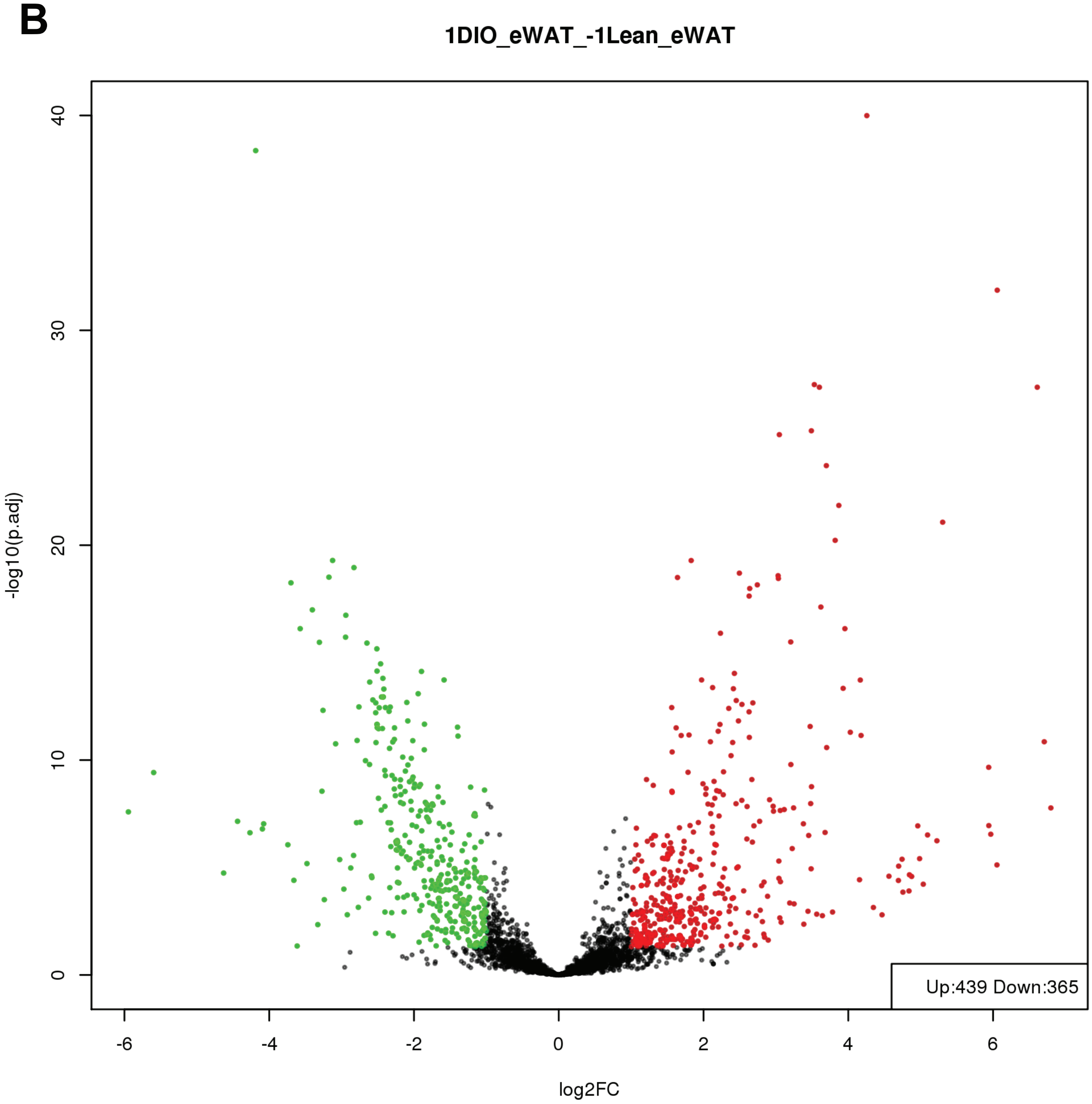
1DIO_eWAT_-1Lean_eWAT
-log10(p.adj)
40
30
20
10
0
-6
-4
-2
0
2
4
6
log2FC
Up:439 Down:365

Fig. S3

C

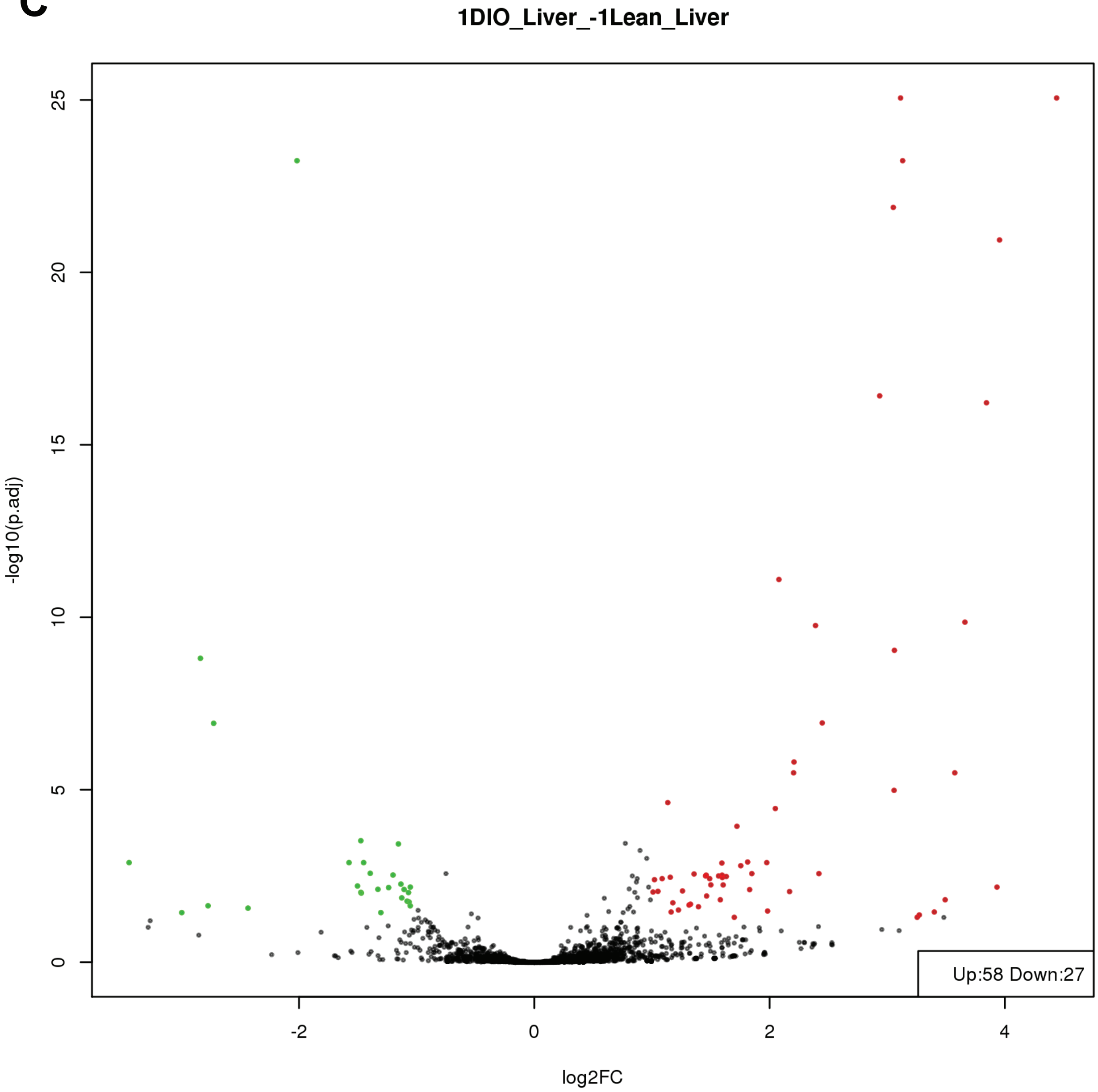

1DIO_Liver_-1Lean_Liver
-log10(p.adj)
0
5
10
15
20
25
-2
0
2
4
log2FC
Up:58 Down:27

Fig. S3

D

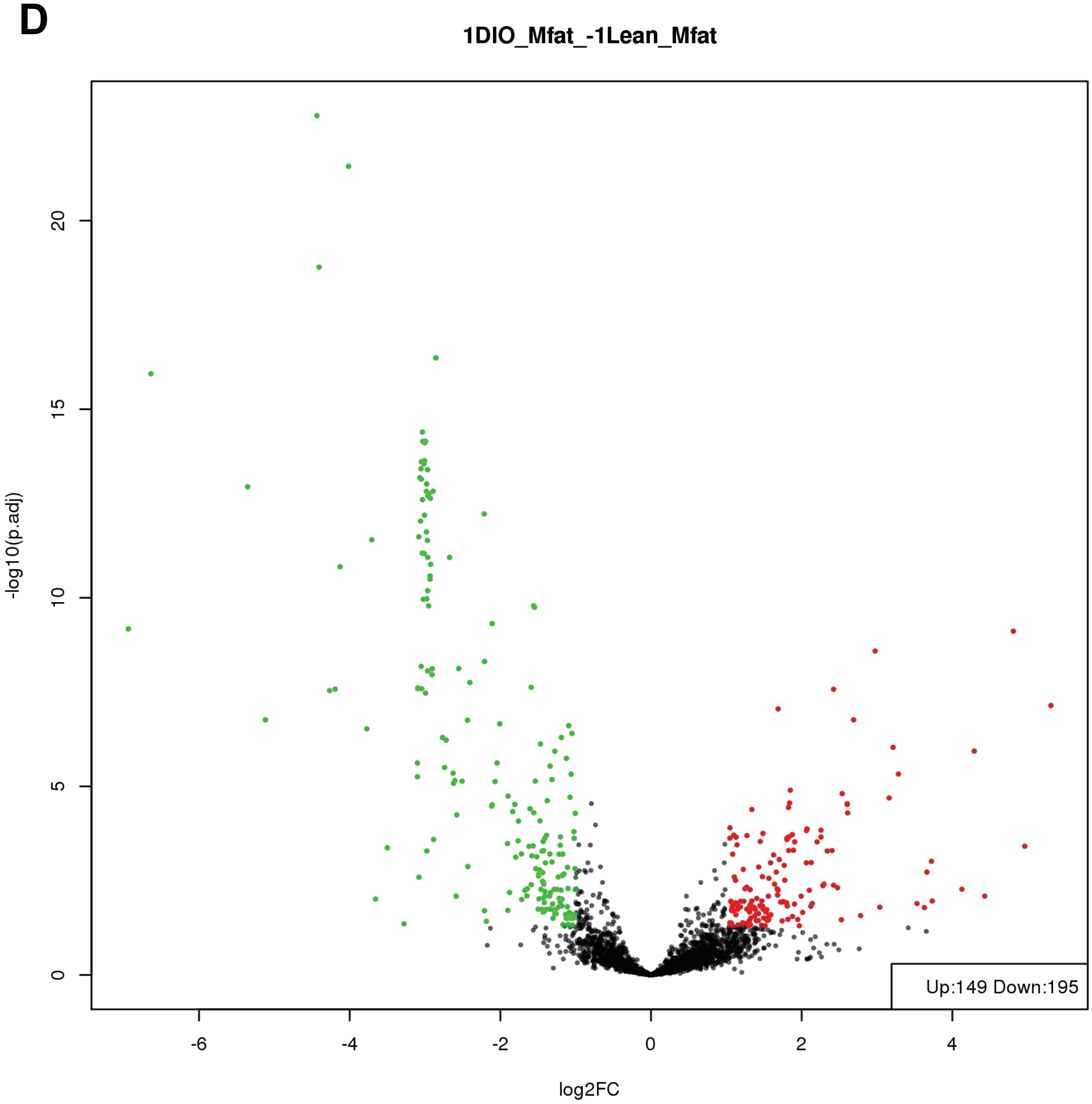
1DIO_Mfat_-1Lean_Mfat
-log10(p.adj)
0
5
10
15
20
-6
-4
-2
0
2
4
log2FC
Up:149 Down:195

### Fig. S3

E

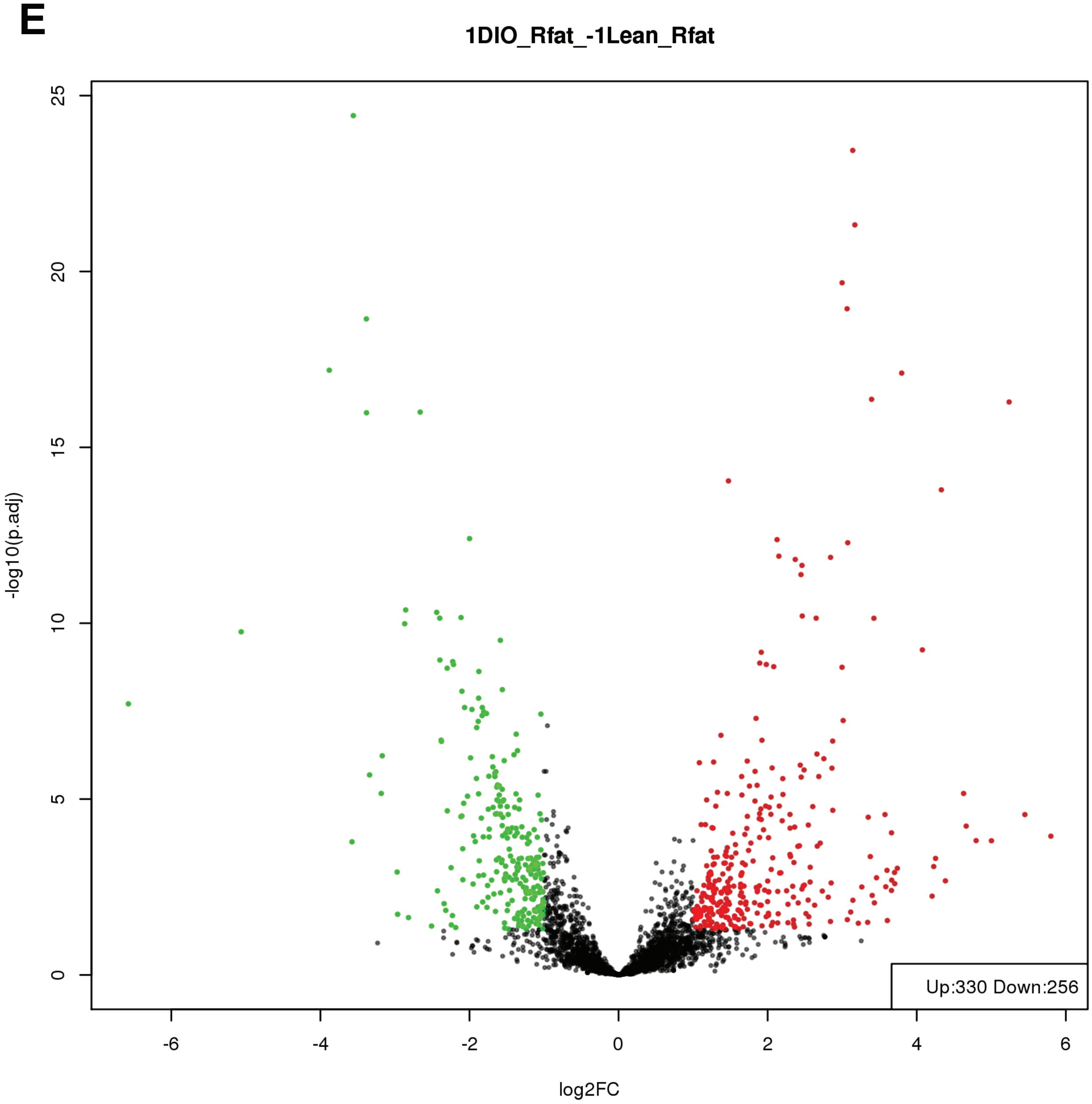
1DIO_Rfat_-1Lean_Rfat
-log10(p.adj)
log2FC
Up:330 Down:256
0
5
10
15
20
25
-6
-4
-2
0
2
4
6

**Fig. S3**

**F**

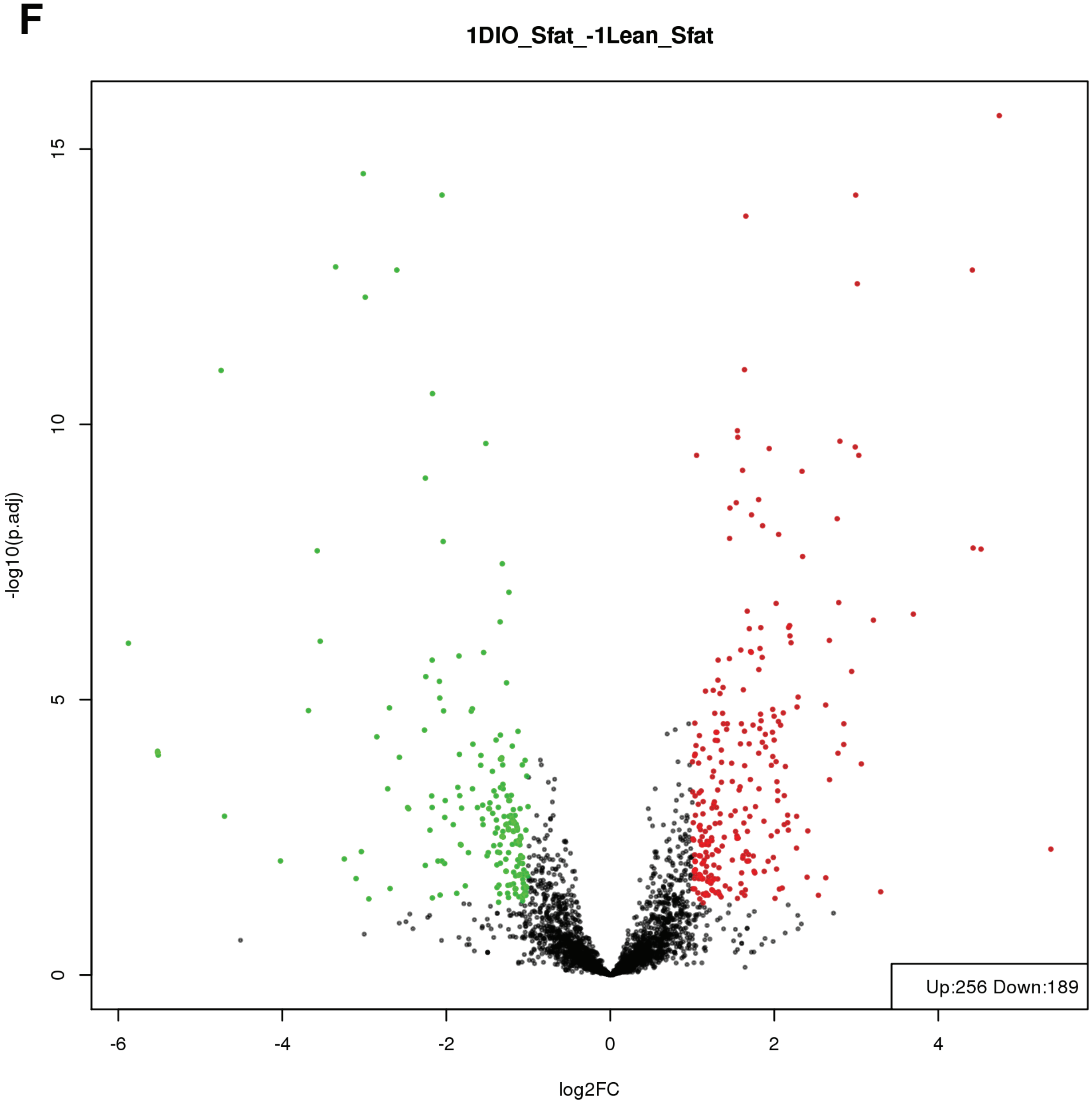

1DIO_Sfat_-1Lean_Sfat
-log10(p.adj)
15
10
5
0
-6
-4
-2
0
2
4
log2FC
Up:256 Down:189

**Fig. S3. Differentially expressed smORF genes in multiple adipose tissue depots and liver using bulk RNA-Seq-based transcriptomics.** smORF coordinates listed in Table S1 were used to extract DIO vs. lean expression levels of all smORF gene coordinates for (A) BAT; (B) epididymal white adipose tissue ("eWAT"); (C) liver; (D) mesentyric adipose tissue ("Mfat"); (E) retroperitoneal adipose tissue ("Rfat"); and (F) subcutaneous adipose tissue ("Sfat"). Changes in RNA expression for adipose protein-coding smORFs induced by DIO in various tissues ($padj < 0.05$ and $|\log2 \text{ fold change}| \geq 1$).

**Fig. S4**

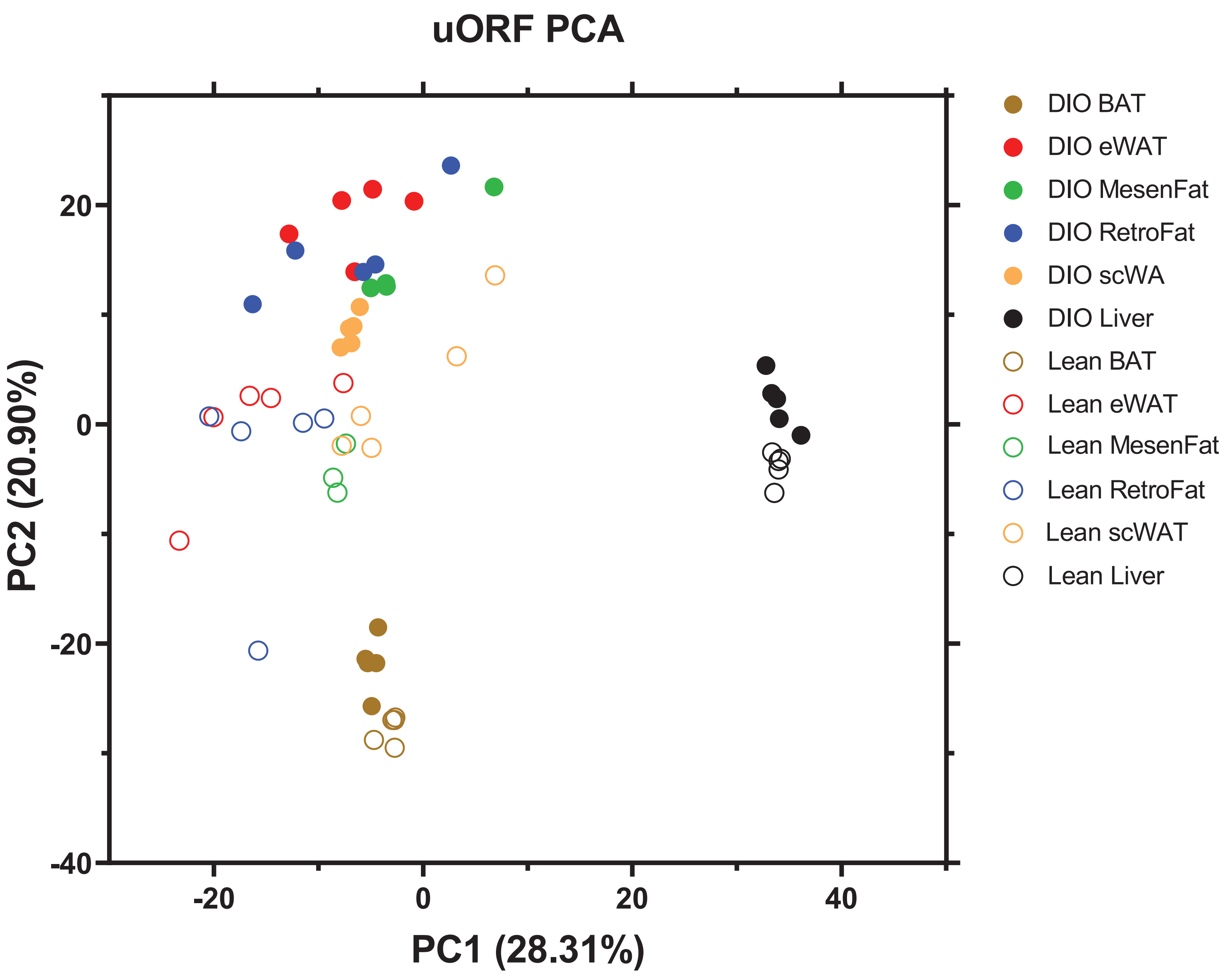
uORF PCA
PC2 (20.90%)
PC1 (28.31%)
20
0
-20
-40
-20
0
20
40
DIO BAT
DIO eWAT
DIO MesenFat
DIO RetroFat
DIO scWA
DIO Liver
Lean BAT
Lean eWAT
Lean MesenFat
Lean RetroFat
Lean scWAT
Lean Liver

**Fig. S4. Adipose protein-coding smORFs are differentially transcribed in diet-induced obesity mice.** Changes in RNA expression for adipose protein-coding smORFs induced by DIO in various tissues (padj $< 0.05$ and $|\log2 \text{ fold change}| \geq 1$) showing PCA analysis of uORF-containing protein-coding smORF RNA expression levels in tissues derived from DIO and lean mice.

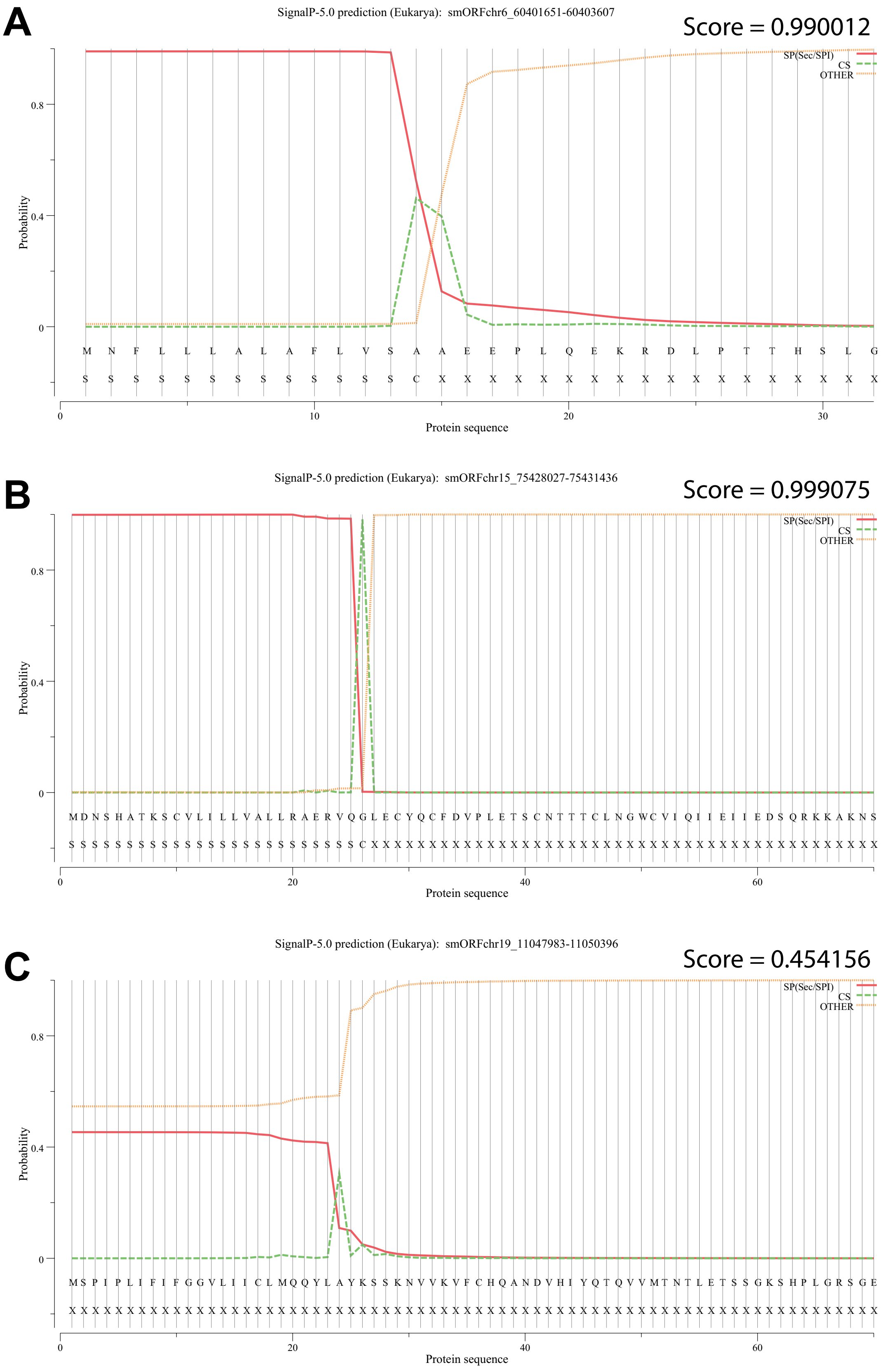
A
SignalP-5.0 prediction (Eukarya): smORFchr6_60401651-60403607
Score = 0.990012
SP(Sec/SPI)
CS
OTHER
Probability
M N F L L L A L A F L V S A A E E P L Q E K R D L P T T H S L G
Protein sequence
B
SignalP-5.0 prediction (Eukarya): smORFchr15_75428027-75431436
Score = 0.999075
SP(Sec/SPI)
CS
OTHER
Probability
M D N S H A T K S C V L I L L V A L L R A E R V Q G L E C Y Q C F D V P L E T S C N T T T C L N G W C V I Q I I E I I E D S Q R K K A K N S
Protein sequence
C
SignalP-5.0 prediction (Eukarya): smORFchr19_11047983-11050396
Score = 0.454156
SP(Sec/SPI)
CS
OTHER
Probability
M S P I P L I F I F G G V L I I C L M Q Q Y L A Y K S S K N V V K V F C H Q A N D V H I Y Q T Q V V M T N T L E T S S G K S H P L G R S G E
Protein sequence

**Fig. S5. Signal peptide predictions of microproteins identified in plasma from DIO/lean young/old mouse experiment.** Microprotein sequences were input into Signal P 5.0 (Almagro Armenteros et al., 2019) . Predictions are depicted above the threshold of 0.4. (A) prediction for microprotein from chr6:60401651-60403607; (B) prediction for microprotein from chr15:75428027-75431436; and (C) prediction for microprotein from chr19:11047983-11050396.

#### SUPPLEMENTARY TABLES

**Table S1: Compilation of all filtered smORFs derived from the Ribo-Seq analysis of primary differentiated brown adipocytes "BAT", primary differentiated beige adipocytes "beige", and primary differentiated white adipocytes "WAT". Table S1 has been filtered such that none of the microprotein sequences appear in the SwissProt database.**

- Sheet 1 ("smORFs_All_SP_Filtered") represents the entire list of smORFs with their chromosomal coordinates, microprotein sequences, and associated genes (where applicable) among other pieces of information for each smORF.
- Sheet 2 ("ALL_smORFs_fasta_formatted") represents all of the smORFs formatted into fasta format that can be amended to a proteome database for mass spectrometry.
- Sheet 3 ("Conserved_smORFs") represents the 204 smORFs with a positive Mean PhyloCSF score indicating conservation.
- Sheet 4 ("tBLASTn_Results") represents the 241 smORFs that have high microprotein amino acid sequence similarity to ORFs found on human transcripts indicating possible conservation between mouse and human.

**Table S2: Signal peptide and transmembrane domain predictions of all smORFs in the Ribo-Seq derived smORF-encoded microprotein library generated in Table S1.**

- Sheet 1 ("SignalP_5.0_predictions") is the output for Signal P 5.0 for signal peptide predictions and
- Sheet 2 ("TMHMM_2.0_predictions") is the output of TMTMM 2.0 for transmembrane domain predictions.
- Sheet 3 ("Phobius_predictions_raw_output") includes all of the unfiltered predictions from Phobius.
- Sheet 4 ("Phobius_signal_pep_positive") includes only the microproteins predicted to have a signal peptide from the Phobius results.
- Sheet 5 ("SignalP_5.0_Phobius_overlap") is the overlap of the SignalP_5.0 and Phobius signal peptide predictions

**Table S3: gene ontology (GO) biological processes analysis on the annotated genes containing uORFs as listed in Table S1.**

**Table S4: compilation of smORF-encoded microproteins identified with mass spectrometry in primary brown, beige, and white adipoccyte cultures using a standard database match search of the canonical uniprot reviewed proteome amended with the microproteins compiled in Table S1.**

- Sheet 1 ("interact.pep-1%FDR_smORFs_only) is the raw search results from Comet + TPP / PeptideProphet filtered for a 1% peptide FDR and reporting only the smORF-encdoed microproteins.
- Sheet 2 ("all_MS-IDed_smORFs_fasta_format") is the summation of unique IDs in Sheet 1 formatted in fasta format for proteomics searches.
- Sheet 3 ("Signal_P_prediction_summary") is the output of Signal P 5.0 with the the fasta file from Sheet 2 as the input.
- Sheet 4 ("smORFs_quantified_w_DIA-MS") is the list of microproteins that were quantifiable in the DIA-MS epxeriment suggesting higher abundance.

**Table S5: Bulk RNA-Seq-based differential expression transcriptomics analysis of all genes in brown adipose tissue (BAT), epididymal white adipose tissue (eWAT), liver, mesentyric adipose tissue (Mfat), retroperitoneal adipose tissue (Rfat), and subcutaneous white adipose tissue (Sfat). Changes in RNA expression for adipose protein-coding smORFs induced by DIO in various tissues (padj < 0.05 and |log2 fold change| ≥ 1).**

**Table S6: Bulk RNA-Seq-based differential expression transcriptomics analysis of all smORF in brown adipose tissue (BAT), epididymal white adipose tissue (eWAT), liver, mesentyric adipose tissue (Mfat), retroperitoneal adipose tissue (Rfat), and subcutaneous white adipose tissue (Sfat). Changes in RNA expression for adipose protein-coding smORFs induced by DIO in various tissues (padj < 0.05 and |log2 fold change| ≥ 1).**

**Table S7: compilation of smORF-encoded microproteins identified with mass spectrometry in the DIO/lean young/old mouse plasma study using a standard database match search of the canonical uniprot reviewed proteome amended with the microproteins compiled in Table S1.**

- Sheet 1 ("interact.pep-1%FDR_smORFs_only) is the raw search results from Comet + TPP / PeptideProphet filtered for a 1% peptide FDR and reporting only the smORF-encdoed microproteins.
- Sheet 2 ("all_MS-IDed_smORFs_fasta_format") is the summation of unique IDs in Sheet 1 formatted in fasta format for proteomics searches.
- Sheet 3 ("Signal_P_prediction_summary") is the output of Signal P 5.0 with the the fasta file from Sheet 2 as the input.

#### RESOURCE AVAILABILITY

##### Lead contact

##### Materials availability

This study did not generate new unique reagents in sufficient quantities to share broadly.

##### Data and code availability

The study did not generate new code. The raw data underlying the figure panels are available in either of the GEO or ProteomeXchange databases (currently proteomics data is stored in the MassIVE database and will become available in the ProteomeXchange upon publication) as listed in the Key Resources Table.
